# AI-Based Comparative Transcriptomics and Gene Network Profiling of *Staphylococcus aureus* in Astronaut-Associated Missions

**DOI:** 10.1101/2025.08.22.671750

**Authors:** Xosé Manuel Tomé Castro

## Abstract

**Introduction:** Spaceflight-associated microgravity alters microbial physiology, raising concerns about the adaptability and pathogenicity of opportunistic bacteria such as *Staphylococcus aureus*. Understanding transcriptomic responses in this context is essential for astronaut health and mission safety.

**Methods:** Publicly available transcriptomic datasets were retrieved from the NASA GeneLab repository and analyzed in two stages. First, three experimental scenarios were compared: (1) BRIC-23 mission (*in vitro* Petri dish cultures aboard the ISS; 9 space vs. 9 ground), (2) SpaceX Inspiration4 mission microbiota (40 samples from 10 body sites of 4 astronauts across flight phases), and (3) Dragon capsule surface cultures (30 samples across 10 capsule zones and 3 time points). Differential expression (DESeq2), functional enrichment, heatmaps, co-expression network analysis (WGCNA), and bootstrapping were applied to identify conserved transcriptomic signatures.

In the second stage, five candidate genes were selected from 45 consistently altered genes across all conditions. These were validated through statistical significance (BRIC-23, *p <* 0.05, significant log_2_ fold change) and consistent presence within co-expression modules. Candidate genes were further integrated with literature-curated virulence and biofilm-associated genes to construct a functional metabolic network using pathway analysis and K-means clustering.

**Results:** Across independent datasets, *S. aureus* displayed convergence in transcriptomic profiles, particularly involving genes linked to virulence, adhesion, biofilm formation, and metabolic adaptation. Network-level integration revealed metabolic reprogramming and transcriptional plasticity as conserved responses, suggesting tightly regulated adaptation rather than random changes.

**Discussion:** The consistent identification of virulence and biofilm-associated genes across heterogeneous datasets indicates that microgravity imposes selective pressure favoring traits that enhance persistence in extreme environments. These findings support the hypothesis that *S. aureus* utilizes adaptive regulatory circuits to balance growth, survival, and pathogenic potential in the spaceflight niche.

**Conclusions:** Our integrative bioinformatics approach reveals conserved adaptive strategies in *S. aureus* under microgravity, characterized by transcriptomic convergence and metabolic reprogramming. These insights underscore the necessity of experimental validation and phenotypic assays to assess microbial risks for astronaut health and to design countermeasures for future long-duration missions.

## Introduction

*Staphylococcus aureus* (*S. aureus*) is a Gram-positive bacterium notable for its extraordinary ability to infect virtually any organ or tissue in the human body. Its pathological spectrum ranges from mild skin infections to potentially life-threatening diseases such as pneumonia, endocarditis, bacteremia, or toxic shock syndrome [41, 43, 22]. As one of the leading causes of hospital-acquired infections, particularly post-surgical infections, its high antibiotic resistance is associated with serious complications in clinical treatment [26].

The vast majority of *Staphylococcus aureus* strains are multi-resistant (MRSA, methicillin-resistant *S. aureus*), making MRSA one of the most lethal bacterial species. Studies underscore the continued public health threat posed by MRSA, which the CDC currently estimates causes more than 70 000 severe infections and approximately 9 000 deaths annually in the United States [9]. Although antimicrobial-resistant hospital-onset infections surged by nearly 20% during the COVID-19 pandemic—peaking in 2021—and remained above pre-pandemic levels in 2022 for most pathogens, MRSA rates notably declined back to pre-pandemic levels by 2022 [8]. Despite this apparent recovery, MRSA continues to impose a substantial burden, particularly in bloodstream infections: 30-day mortality ranges from about 6.9% in individuals younger than 65 to over 33% in those older than 80. During the pandemic, overall mortality from *S. aureus* bacteremia reached 41.9%, with the highest rates observed among patients older than 60 [37]. To better guide interventions, the CDC plans to update antimicrobial resistance burden estimates, including MRSA, on a biennial basis beginning in 2025 [9, 8].

Although MRSA has traditionally been associated with hospital settings, community-associated strains (CA-MRSA) have emerged, which are often more virulent and transmissible [41, 22]. A particularly concerning example is MRSA252, a variant capable of causing both hospital-acquired and community-acquired infections. This strain belongs to a genetic lineage linked to hospital outbreaks and carries multiple resistance mechanisms, including beta-lactam resistance [26].

It is estimated that approximately 30% of the global population is colonized by *S. aureus*, mainly in the nasal passages and skin, which significantly increases the risk of developing more severe infections [16]. Its pathogenicity is due to a true arsenal of molecular tools that allow it to effectively interact with both biotic and abiotic environments. Among these are surface adhesins for attachment to tissues and medical devices; toxins that enable immune evasion; enzymes that degrade the extracellular matrix, facilitating tissue invasion; and a remarkable ability to form biofilms [27, 47].

Biofilm formation is crucial to understanding its virulence. These microbial communities are protected by a polysaccharide matrix that greatly hinders immune system action and antibiotic penetration, such as vancomycin or methicillin, thereby promoting resistance development [41]. The regulation of biofilm formation involves complex systems such as quorum sensing (mediated by autoinducers), second messengers (c-di-GMP), two-component systems, and non-coding RNAs, which coordinate the transition between planktonic life and the stationary biofilm form [27].

Given its extraordinary adaptability, *S. aureus* represents a critical risk in enclosed and extreme environments, such as the International Space Station (ISS) or future interplanetary missions [10, 25, 16, 30, 29, 44, 10]. Factors such as microgravity, space radiation, and immunosuppressive stress experienced by astronauts significantly amplify this threat, as these biofilms can adhere to spacecraft surfaces, making eradication difficult [30]. In fact, large amounts of *S. aureus* have been isolated from astronauts’ nasal passages after the Apollo missions (13–15) [7], and studies conducted on the ISS confirm that staphylococci dominate the crew’s microbiota as the most prevalent bacteria on surfaces [44]. This situation has motivated extensive research into *S. aureus* virulence and resistance under microgravity stress conditions [42].

On the ISS, *S. aureus* is widely distributed on both equipment and surfaces, as well as on the astronauts themselves. Its presence in these confined environments and its potential for resistance development represent a significant health threat during prolonged missions to the Moon or Mars. Reports on medications used aboard the ISS indicate that 25% of astronauts have required treatment for skin eruptions with antifungals, corticosteroids, or antihistamines [48]. While most treatments were effective, some required medication changes, suggesting limitations in currently available therapies [48]. This issue is exacerbated by space-induced immune alterations and changes in the skin microbiome due to restrictive hygiene methods [33].

*Staphylococcus* species isolated on the ISS show resistance to antibiotics used onboard and a marked ability to form biofilms [33]. Studies, such as the SpaceX-3 mission, detected higher mutation rates in *S. epidermidis* under microgravity, while analyses of surfaces and air confirmed the presence of high-risk microorganisms resistant to *β*-lactams and vancomycin [11].

NASA’s Biological Research in Canisters-23 (BRIC-23) experiment specifically analyzed *S. aureus* and *B. subtilis* in microgravity. Integration of RNA-Seq, proteomics, and metabolomics data revealed in *S. aureus*: increased amino acid metabolism and Krebs cycle activity, decreased glycolysis, and overexpression of *agr* system genes (virulence regulator). The secretome showed higher abundance of *agr* -regulated proteins, suggesting that the space environment profoundly alters its metabolism, enhances quorum sensing, and modulates virulence factors [16].

Space research faces significant methodological challenges, such as small sample sizes (the ISS hosts a maximum of six astronauts) and a lack of standardization between missions. Omics studies are particularly scarce: NASA’s “Twin Study” included only one astronaut in space and their twin on Earth [12], while the Inspiration4 (SpaceX) mission generated multi-omic data from four crew members [31]. On Inspiration4, biological samples were collected at 11 time points (from L-92 to R+194 days), covering a total period of 289 days (2021–2022) [31].

Using mixed models, the nasal and oral microbiota were analyzed, identifying six temporal patterns of change. Tools such as MetaPhlAn and StrainPhlAn tracked strain exchanges between crew members and the environment, while lasso regressions associated microbial genes with immune expression. Results highlighted genes persistently altered post-flight, revealing microbial and immunological adaptations specific to microgravity [44].

These findings underscore the urgent need to develop longitudinal, integrative approaches combining multi-omic data to understand the complex interactions between the human microbiome and the space environment. Such understanding is fundamental to ensuring astronaut health on prolonged missions.

## Results

### Comparative Analysis of RNA-Seq Data

The dimensionality reduction analysis using UMAP, as shown in Figure 1, revealed differentiated patterns in sample distribution according to the experimental scenario studied. In the BRIC-23 dataset, a clear separation between spaceflight and ground control conditions was evident, suggesting a differentiated and robust transcriptomic response to simulated microgravity conditions. In contrast, scenarios corresponding to microbiota (astronauts ASTRO_1 to ASTRO_4) and surface showed no clear or statistically significant separation between different experimental conditions (L-3, FD2, FD3, and R+1), pointing toward greater heterogeneity in response or a smaller magnitude effect of spatial factors in these more complex environments.

**Figure 1:**
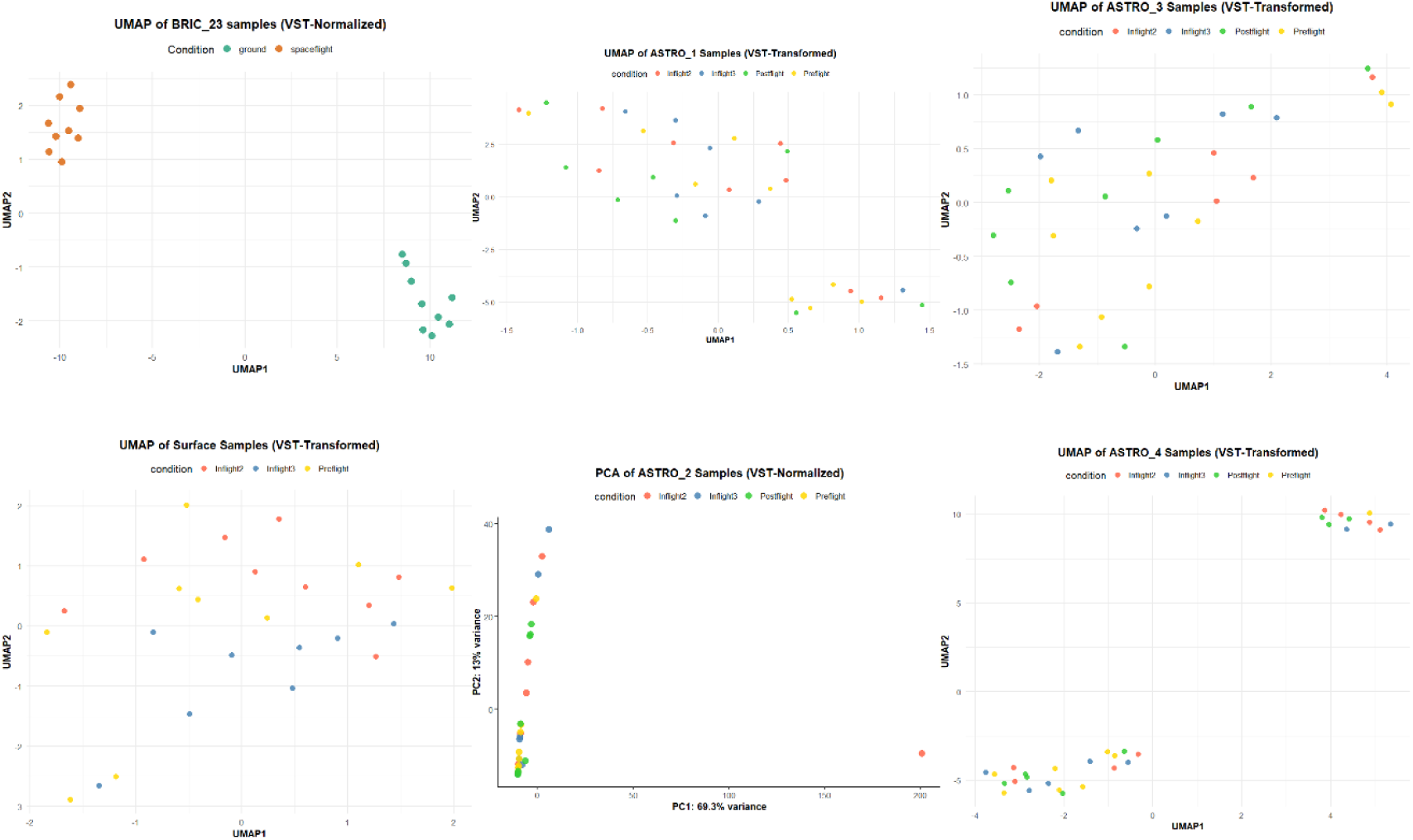
UMAP dimensionality reduction plots for different study scenarios.

A heatmap was constructed, Figure 2, focusing on genes that showed the greatest variability in their expression in BRIC_23, excluding those not expressed in microbiota and surface scenarios, which are included in the document appendix. The heatmap with the most variable genes was chosen as a consequence of the absence of clear and statistically significant separation observed in different scenarios, which made it difficult to extract conclusive information through differentially expressed gene analysis. The selection of most variable genes allowed identification of transcriptional regulation patterns even in the absence of evident differentiation between conditions.

**Figure 2:**
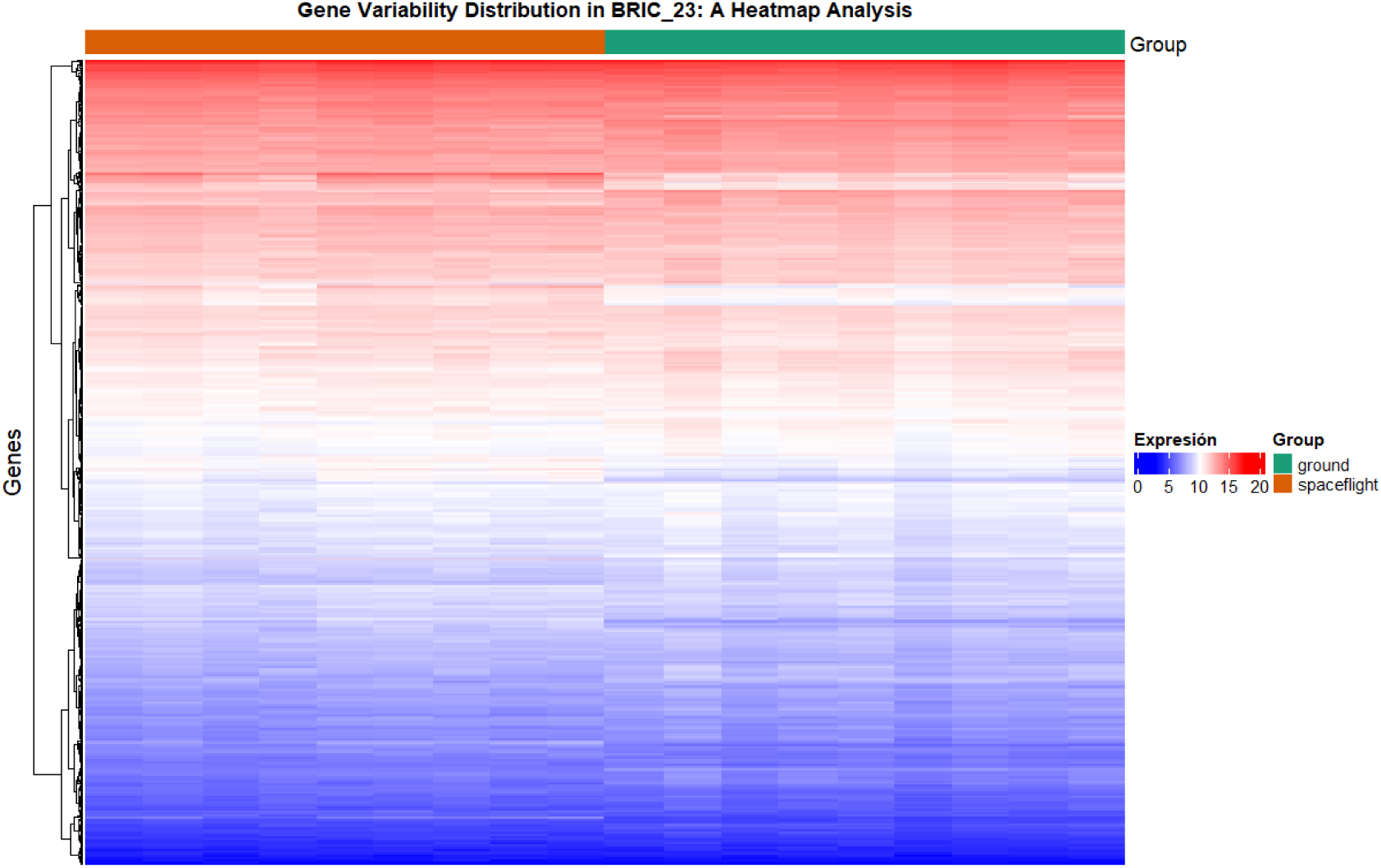
General heatmap of expression variability data by BRIC_23 scenario

In the BRIC_23 dataset, the heatmap revealed a marked distinction between highly expressed genes and genes with low expression, involving a considerable number of genes, thus confirming a solid transcriptomic response to space conditions.

In the WGCNA analysis framework, the correlation between gene co-expression modules and experimental conditions (traits) is evaluated using eigengenes, which represent the characteristic average expression pattern of each module. By correlating these eigengenes with experimental conditions, a biological association metric between the module and the studied trait is obtained, shown in:

- Microbiota 1: 10 functional modules
- Microbiota 2: 4 functional modules
- Microbiota 3: 31 functional modules
- Microbiota 4: 3 functional modules
- Petri: 2 functional modules
- Surface: 27 functional modules

A positive and statistically significant correlation (*p <* 0.05) indicates that the module is more activated or expressed under that specific condition, while a significant negative correlation suggests that the module is repressed or shows lower expression in that condition. Consequently, modules that show high correlations and low p-values are considered biologically relevant and can be prioritized for subsequent functional studies or for identifying key genes (hub genes) involved in the specific treatment response. This information is typically represented through a heatmap, where color intensity and tone reflect the magnitude and direction of correlation, facilitating comprehensive interpretation of the transcriptomic system in relation to the analyzed traits [23].

In the BRIC_23 dataset, only two modules were identified, which showed strong and statistically significant correlations with experimental groups, shown in Figure 3. Specifically, a positive correlation of 0.93 and negative of −0.93 with a p-value of 2 × 10*^−^*^8^ was observed, evidencing a robust and clear association between the identified modules and experimental conditions. This finding suggests that genes grouped in these modules experience differential regulation according to experimental group and, therefore, may be relevant for differentiating biological processes between studied conditions.

**Figure 3:**
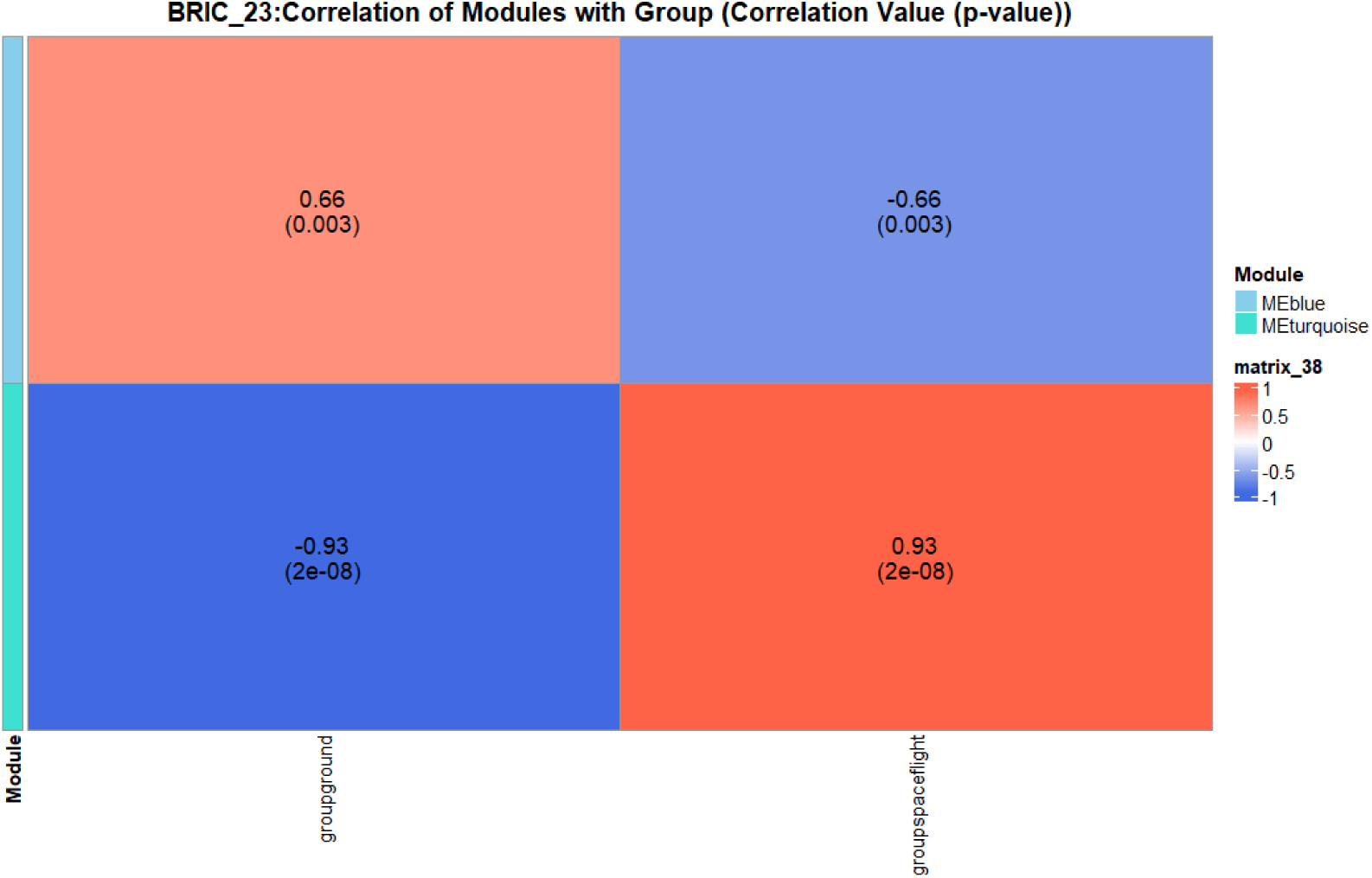
Correlation of gene modules from BRIC_23 dataset with experimental groups. Two modules with significant correlations (0.93 and −0.93; *p* = 2 × 10*^−^*^8^) are highlighted, indicating differential regulation according to condition.

In the ASTRO_1 and ASTRO_3 datasets, as shown in Figure 4, multiple modules were identified, some of which showed moderate correlations (values between 0.03 and 0.05) with acceptable statistical significance. Although not all modules reached statistical significance (*p <* 0.05), interesting patterns were identified that justify subsequent functional analysis, especially in ASTRO_3, where correlations were more notable and consistent.

**Figure 4:**
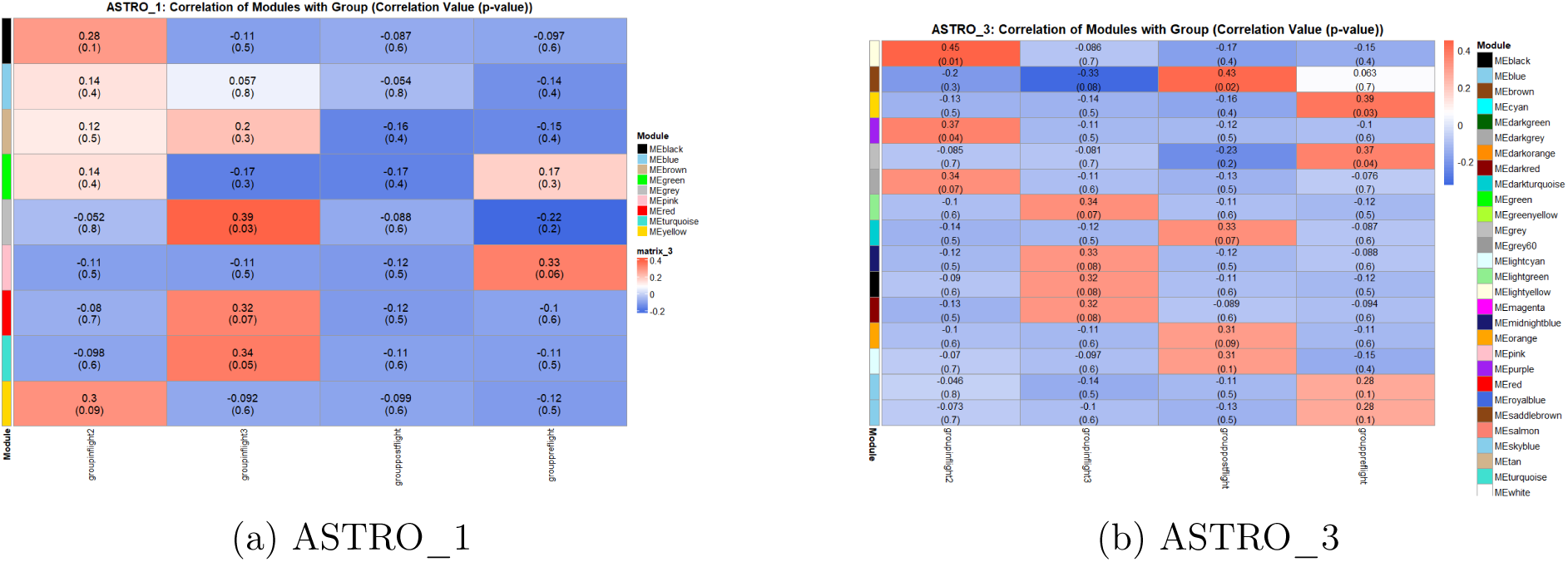
Modules identified in ASTRO_1 and ASTRO_3 datasets. Multiple modules were observed, some with moderate correlations (0.3–0.5) and acceptable statistical significance. In ASTRO_3, patterns were more notable and consistent, justifying subsequent functional analysis.

The ASTRO_4 and Surface datasets showed important limitations in the analysis, evidenced in Figure 5. In ASTRO_4, despite identifying four modules, observed correlations were weak and lacked statistical significance, indicating low robustness in the association between modules and experimental conditions. In the case of Surface, although two modules were identified, it was not possible to establish a clear functional association between genes and predominant biological functions due to data insufficiency. This limitation made it impossible to simultaneously evaluate the most common functions and subcellular localization of grouped genes.

**Figure 5:**
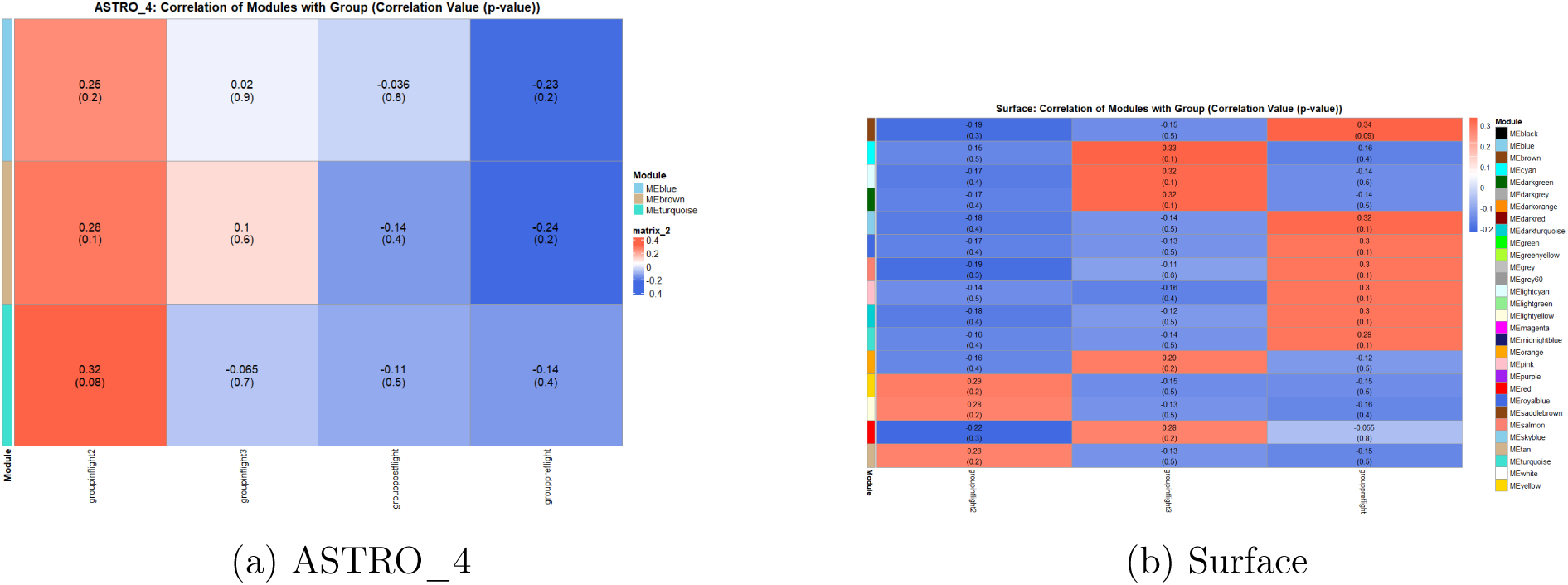
ASTRO_4 and Surface datasets showed important limitations in the analysis. In ASTRO_4, despite identifying four modules, observed correlations were weak and lacked statistical significance, indicating low robustness in the association between modules and experimental conditions. In the case of Surface, although two modules were identified, it was not possible to establish a clear functional association between genes and predominant biological functions due to data insufficiency, making it impossible to simultaneously evaluate the most common functions and subcellular localization of grouped genes.

It is important to note that in microbiota 1 and 3, only one robust module was obtained in each case, which limited comparative analysis but allowed identification of at least one coherent gene set associated with some experimental condition. In contrast, microbiota 2 and 4, similar to BRIC_23, showed greater modular richness, facilitating more detailed and comprehensive functional interpretation.

Module stability analysis through bootstrapping constitutes a critical stage in WGCNA for validating the robustness of identified co-expression modules. This methodological approach allows distinguishing between robust co-expression patterns and those that could arise from random sampling variations, thus providing a quantitative measure of the reliability of gene groupings.

The interpretation of these results should consider that red blocks on the heatmap’s main diagonal confirm intramodular stability, while low-intensity regions indicate inconsistencies in gene assignment. This information is fundamental for selecting reliable modules for subsequent functional analyses, since only those demonstrating high stability provide a solid foundation for biological inferences.

The stability heatmaps obtained in Figure 6 reveal substantial differences in modular robustness between evaluated experimental conditions. In BRIC_23, ASTRO_1, ASTRO_2, and ASTRO_4, clearly delineated blocks with intense red coloration (stability values near 1.0) are observed, indicating that genes group consistently in the same module through multiple resampling iterations. This consistency suggests the presence of highly stable modules that reflect robust underlying biological patterns.

**Figure 6:**
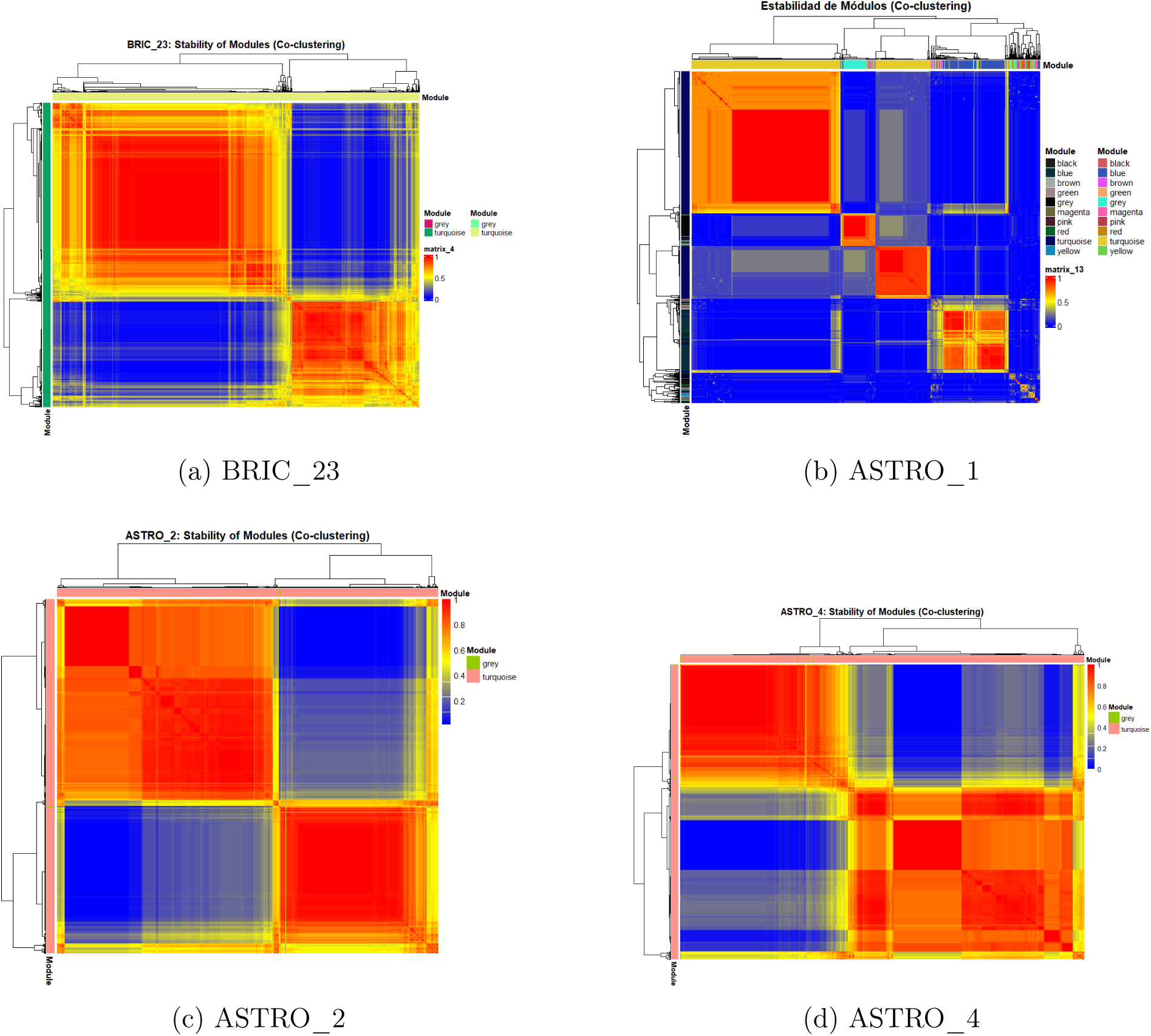
Modular stability heatmaps for BRIC_23, ASTRO_1, ASTRO_2, and ASTRO_4. Clearly defined red blocks (values near 1.0) indicate that genes group consistently in the same modules after multiple resampling, suggesting highly stable modules and robust biological patterns.

On the other hand, Surface and ASTRO_3 (Figure 7) exhibit markedly different patterns, characterized by diffuse regions or bluish tonality (values close to 0.0). This coloration indicates that genes do not maintain consistent groupings during the bootstrapping process, suggesting unstable modules or genes without clear modular assignment. The absence of well-defined blocks in these cases signals that initially identified patterns might not be reproducible under resampling conditions.

**Figure 7:**
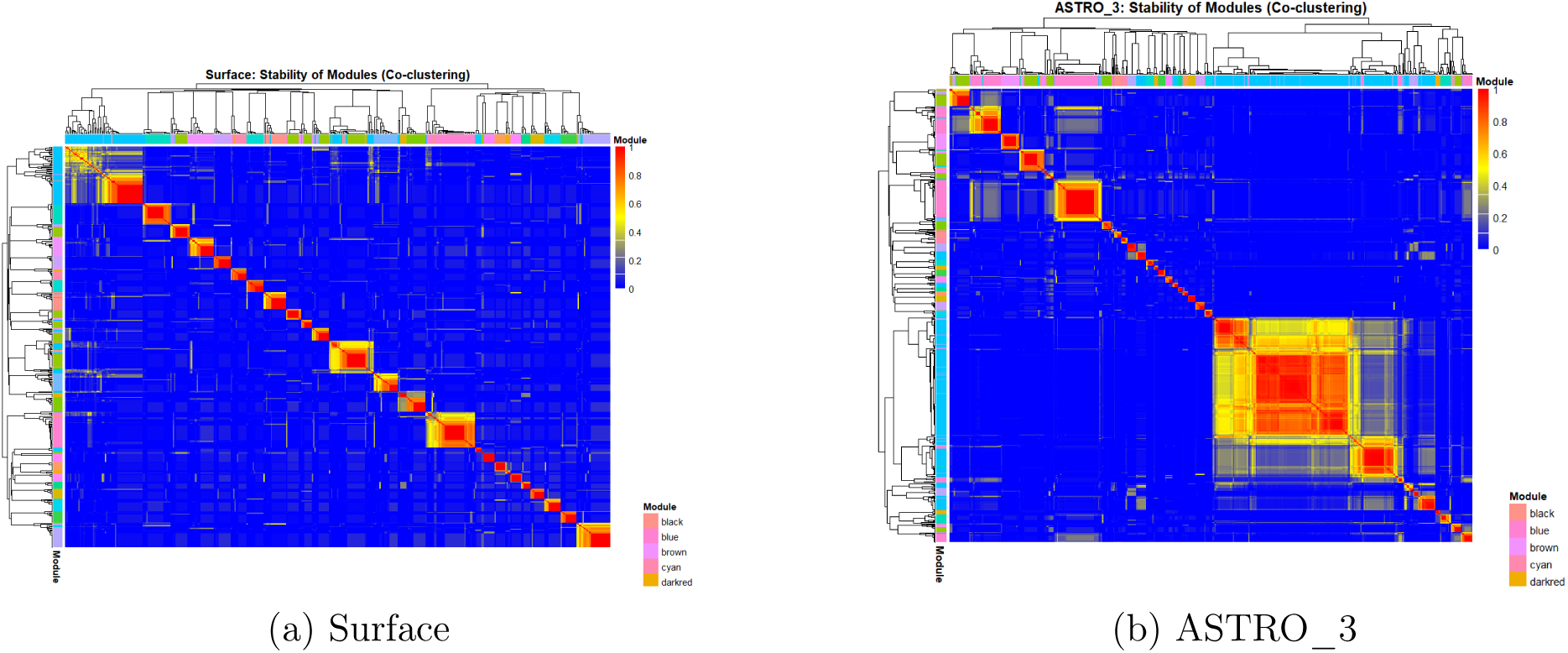
Modular stability heatmaps for Surface and ASTRO_3 datasets. Diffuse patterns or bluish tonality (values close to 0.0) are observed, indicating that genes do not maintain consistent groupings during bootstrapping, suggesting unstable modules or genes without clear modular assignment. The absence of well-defined blocks signals that initially identified patterns might not be reproducible under resampling.

Following bootstrapping analysis, functional enrichment of modules that demonstrated greatest statistical robustness was performed. WGCNA analysis results revealed heterogeneous distribution in the number of functional modules detected in each experimental scenario

The obtained results revealed significant differences in distribution and functional characteristics between different evaluated experimental scenarios.

The analysis revealed a heterogeneous distribution of robust modules among different experimental scenarios. ASTRO_1 and ASTRO_3 scenarios presented one robust module each, while ASTRO_2, ASTRO_4, and BRIC_23 exhibited four, three and two robust modules respectively. The Surface scenario showed two modules, although limitations in data availability prevented performing a complete detailed functional association for both modules simultaneously. The functions of each scenario along with the most relevant genes can be observed in Table 1.

**Table 1:** Summary of Functional Enrichment of Robust Modules by Scenario.

| Scenario | Main Functions | Relevant Genes |
| --- | --- | --- |
| ASTRO_1 | Cellular stress response; chromosome segregation; intracellular pH regulation; virulence control | *GrpE*, *DnaJ*, *DnaK*, *XerD*, *XerC*, *MnhA1/D1*, *SaeR/SaeS* |
| ASTRO_2 | Essential metabolic processes; ribosomal assembly; sugar transport and phosphorylation (PTS) | *RplC*, *Rps*, *Pdh*, *crr/mtlA* (LACe) |
| ASTRO_3 | Essential metabolic processes; ribosomal assembly; energy metabolism | *RplC*, *Rps*, *Pdh* |
| ASTRO_4 | Ribosomal and respiratory functions; carbon metabolism; peptidoglycan synthesis | *RplC*, *Rps*, *qoxA*, *crr/mtlA*, *gcvT*, *murA2* |
| BRIC-23 | Energy metabolism; ABC and nickel transport; hemolytic and leukotoxic activity | *RplC*, *Rps*, *Pdh*, *mtlF*, *nik*, *hlgA*, *hlgB*, *hlgC* |
| Surface | Complement inhibition; bacterial adhesion; amino acid biosynthesis | *Sbi*, *ilvD*, *clfA*, *sodM* |

**Table 2:** Genes Detected Across Experimental Scenarios on Petri Plates and Microbiota Conditions.

| Gene | Experimental Scenario(s) |
| --- | --- |
| *mco*<br>(gene-SAR_RS03660) | Microbiota 2, Microbiota 4, Petri |
| *gluD*<br>(gene-SAR_RS04685) | Microbiota 1, Microbiota 2, Microbiota 4, Petri |
| *odhA*<br>(gene-SAR_RS07275) | Microbiota 1, Microbiota 4, Petri |
| *vraR*<br>(gene-SAR_RS10305) | Microbiota 2, Microbiota 4, Petri |
| *argF*<br>(gene-SAR_RS14275) | Microbiota 2, Microbiota 4, Petri |

To understand the distribution of biological functions and relevant genes in analyzed scenarios, two key graphical representations were used.

First, a function heatmap (Figure 8) illustrated the presence or absence of main functions in each scenario. Color intensity indicated activation or inactivity of each function. This analysis revealed that energy metabolism, ribosomal and respiratory functions were widely distributed (ASTRO_2, ASTRO_3, ASTRO_4, and BRIC-23). In contrast, sugar transport was concentrated in ASTRO_1, ASTRO_2, and ASTRO_4. Specialized functions, such as cellular stress response, were exclusive to ASTRO_1, while virulence and bacterial adhesion were observed specifically in BRIC-23 and Surface, respectively.

**Figure 8:**
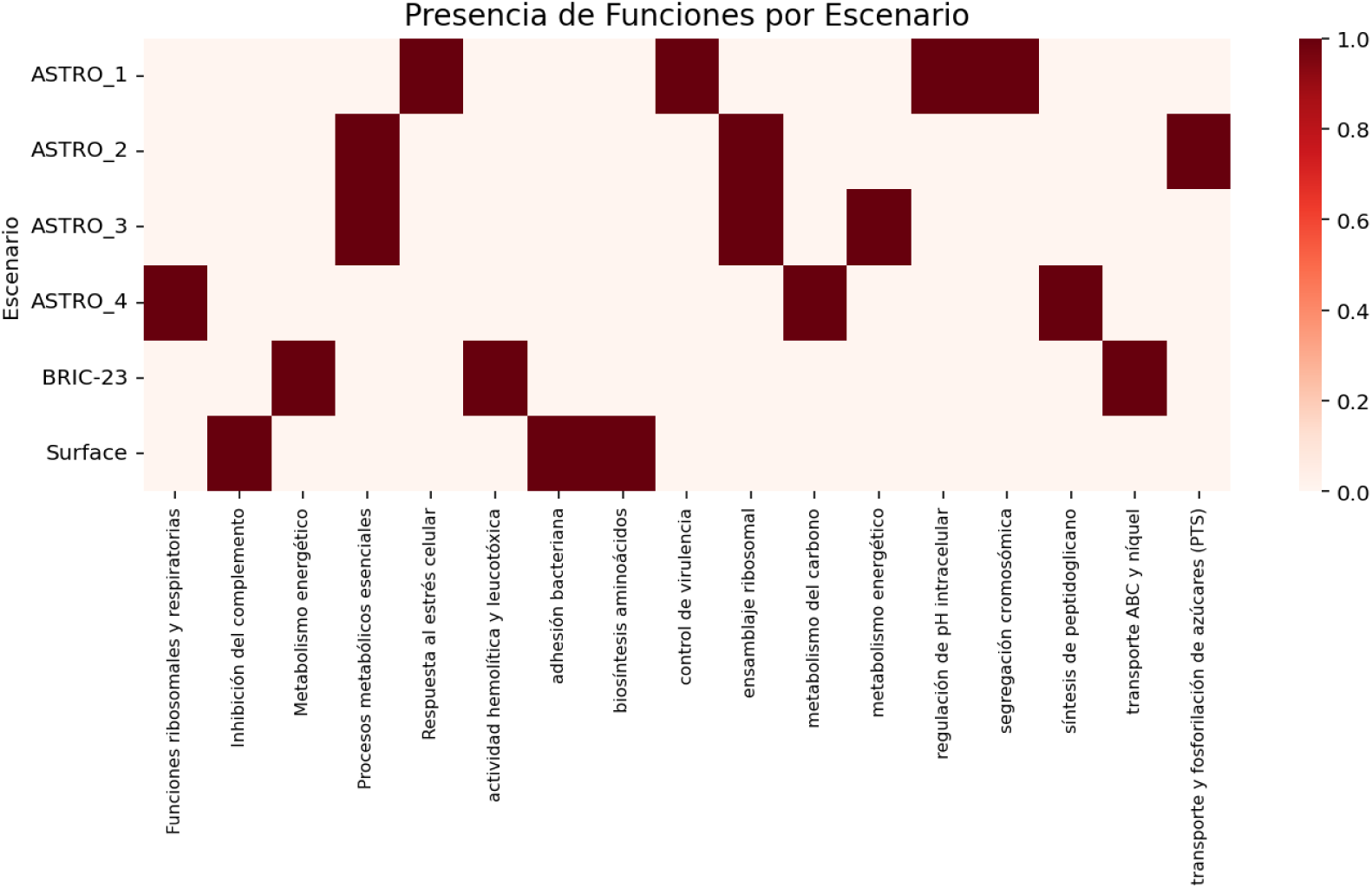
Heatmap showing the presence or absence of main biological functions in different experimental scenarios. Color intensity reflects activation or inactivity of each function, allowing identification of shared and specific functional patterns that contribute to transcriptomic adaptation of *Staphylococcus aureus* under space conditions.

Second, a bar chart of most shared genes (Figure 9) highlighted genes present in multiple scenarios. The genes *RplC* and *Rps*, crucial for ribosomal protein synthesis, showed the greatest distribution, appearing in four scenarios (ASTRO_2, ASTRO_3, ASTRO_4, and BRIC-23). The *Pdh* gene, linked to energy metabolism, was found in three scenarios (ASTRO_2, ASTRO_3, and BRIC-23). Finally, sugar transport system genes *crr/mtlA* were present in two scenarios (ASTRO_2 and ASTRO_4), suggesting their relevant role in active metabolite transport.

**Figure 9:**
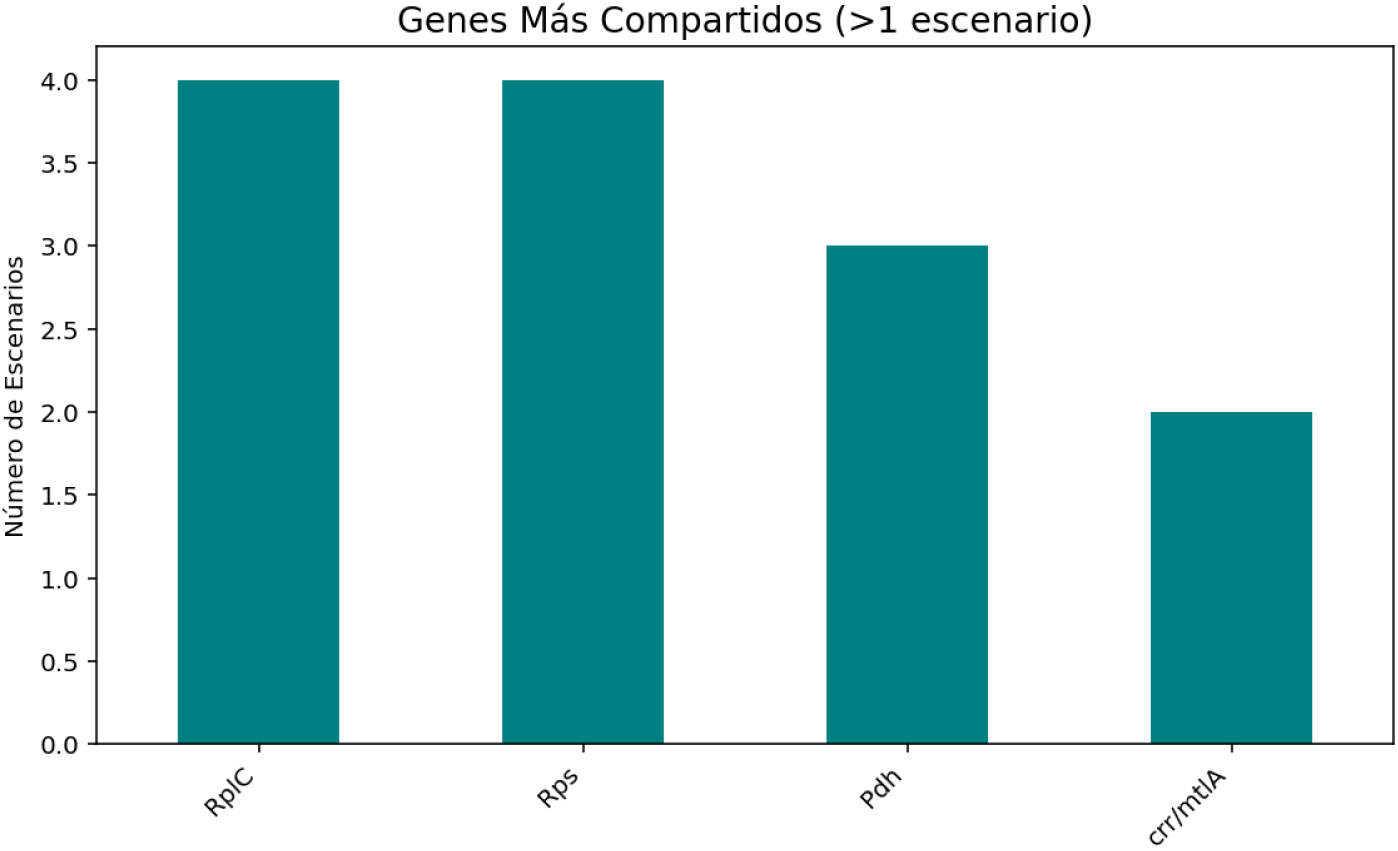
Bar chart representing the most shared genes among different scenarios. It highlights genes like *RplC* and *Rps*, essential for ribosomal protein synthesis, and *Pdh*, involved in energy metabolism, showing their presence in multiple scenarios and their probable central role in bacterial adaptation.

### Synthetic Biology-Based Transcriptomic Analysis of Common Genes in Modules

From the first data set, a total of **45 unique genes** were identified. These genes were present in all scenarios (BRIC_23, astro1-4, and surface) and across all conditions evaluated within each one. This information was preserved in a file called genes_comunes_all_entre_prueba facilitating its accessibility for subsequent analyses. In the second set of analyses, where *log2 fold change* and *p-value* were used, the only scenario that yielded statistically significant genes was BRIC_23. Although the same procedure was applied to the other experimental scenarios, no statistically significant genes were detected. Therefore, the Volcano Plot shown in Figure 10, corresponding to the BRIC_23 scenario, was used, as it was the only one that demonstrated statistical significance. The analysis focused on this scenario because it could provide more statistically relevant results; however, it should be noted that it does not reflect whether genes are upregulated or downregulated in microbiota or surface scenarios. This is because it does not represent actual bacterial competition with other species, but rather their behavior under isolated pure culture conditions on Petri plates.

**Figure 10:**
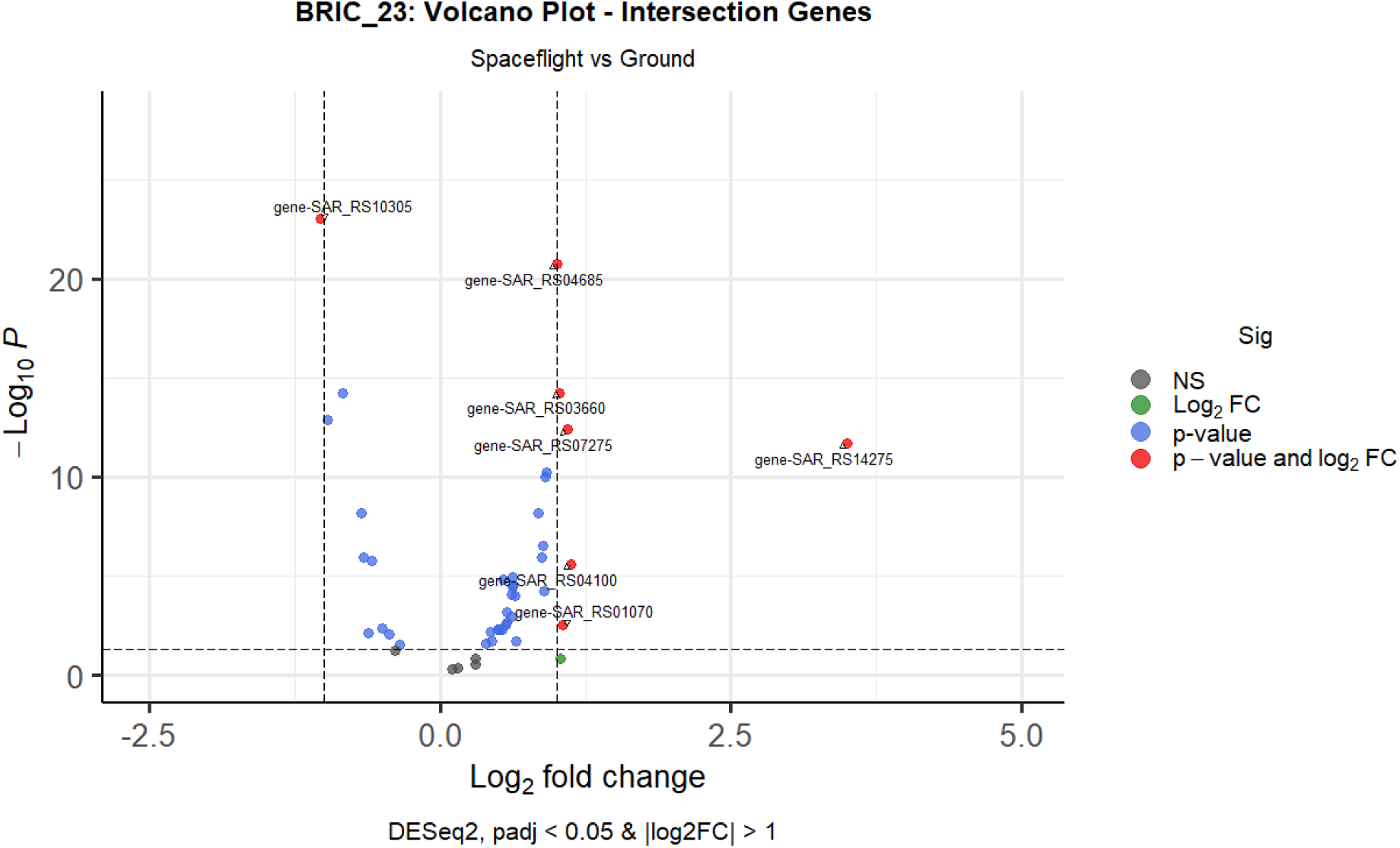
Volcano Plot of BRIC_23 scenario showing the 45 analyzed genes.

In the obtained Volcano Plot, it was observed that of the 45 analyzed genes, only seven were statistically significant when applying the established thresholds of log_2_ fold change and *p*-value: *vraR* (gene-SAR_RS10305), *pflB* (gene-SAR_RS01070), *hpf* (gene-SAR_RS04100), *odhA* (gene-SAR_RS07275), *argF* (gene-SAR_RS14275), *mco* (gene-SAR_RS03660), and *gluD* (gene-SAR_RS04685).

From these 7 genes, the third set was performed to verify their presence in selected modules after bootstrap with WGCNA. As a result of this analysis, only 5 genes were retained in modules considered most stable, reproducible, and robust. These five genes constitute the most promising biological candidates identified in our analysis, by simultaneously integrating functional, statistical, modular, and experimental robustness criteria, thus providing a solid foundation for future research and experimental validations. Genes that passed this selection filter, along with detailed documentation of conditions in which they were detected, were exported to a CSV file named genes_petri_condiciones_min_3.csv, facilitating their accessibility for subsequent analyses and experimental validation studies shown in the following table summarizes the experimental scenarios in which the five selected genes were detected. Scenarios include various microbiota conditions (Microbiota 1–4) and isolated Petri plate cultures, showing the consistency of gene detection across multiple conditions.

The K-means clustering algorithm (with *k* = 2) was implemented (Figure 11) to group genes according to their expression profiles, with the objective of identifying coherent patterns in different functional pathways. This procedure resulted in assigning each gene to one of two clusters (0 or 1), representing groups with similar pathway profiles. Cluster 0 comprises genes whose pathways show no functional relationship among themselves, while Cluster 1 groups those genes that share similar metabolic pathways.

**Figure 11:**
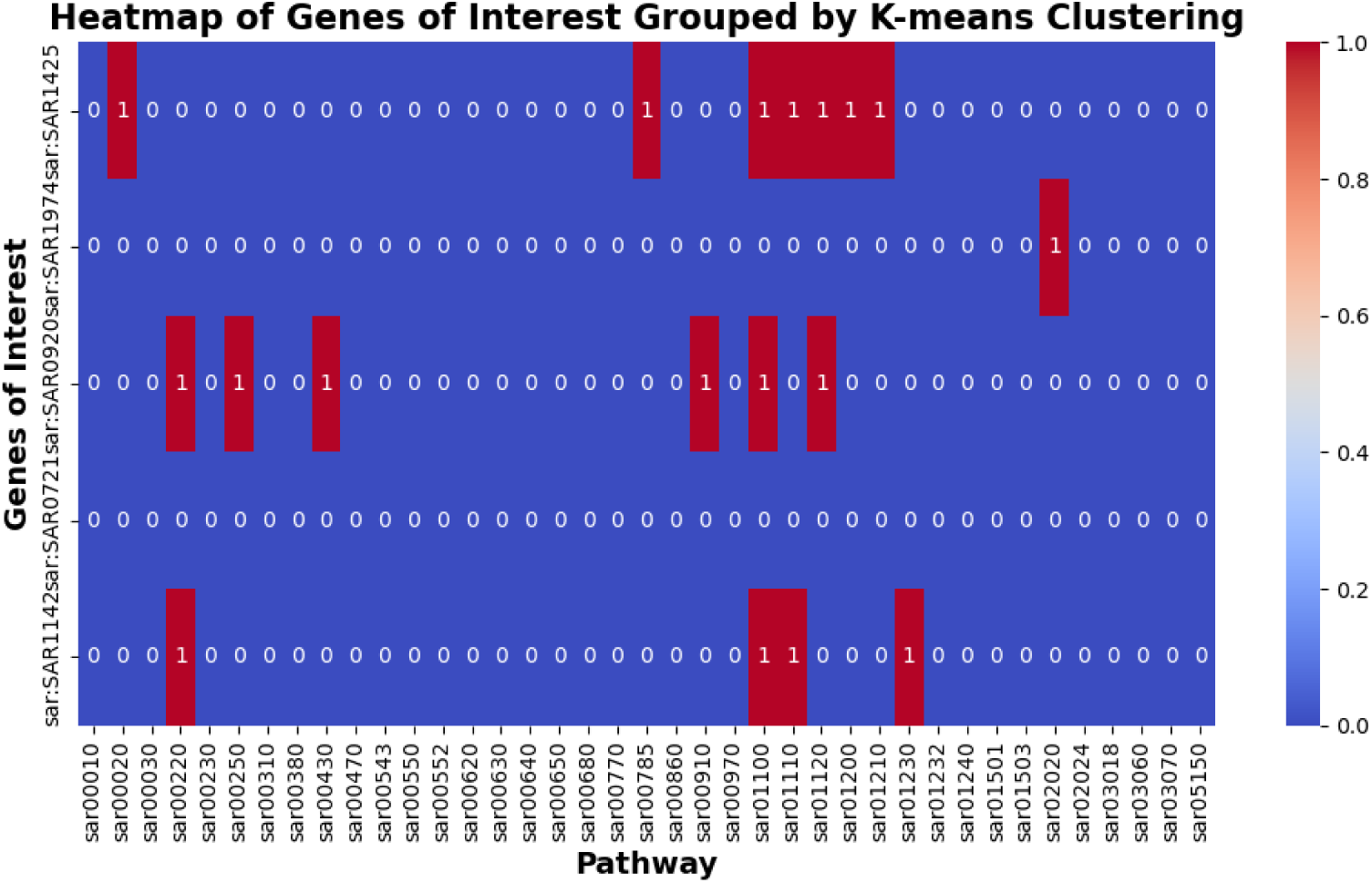
Gene grouping through K-means clustering with *k* = 2. Gene assignment to two distinct clusters based on their expression profiles is observed. Cluster 0 contains genes whose functional pathways are unrelated, while Cluster 1 groups genes with similar metabolic pathways.

Although this classification is not based on predefined biological labels, it allows identification of functional modules or gene co-expression that could be associated with common biological processes. Data visualization through a heatmap revealed marked differences in expression between both groups, highlighting that the gene *sar:SAR0721* shows no common pathway with other analyzed genes.

After gene grouping through K-means clustering (with *k* = 2), gene expression patterns were explored to identify biologically significant groupings. To facilitate their interpretation, specialized dimensionality reduction techniques were applied.

UMAP analysis revealed clear separation between clusters 0 and 1. Together, these analyses demonstrate that genes present expression patterns that allow their coherent grouping, opening new perspectives for interpreting their common functions, performing functional enrichment studies, and guiding future experimental investigations.

A metabolic pathway network was constructed that interconnects genes and pathways using previously obtained binary data about their functional association, shown in Figure 12. From the expression DataFrame, a graph was generated using NetworkX where nodes represent genes and pathways, and edges indicate their functional relationship. Genes of interest were highlighted to emphasize their role in the network and both pathways associated with these genes and other genes sharing these pathways were extracted, thus expanding the functional context.

**Figure 12:**
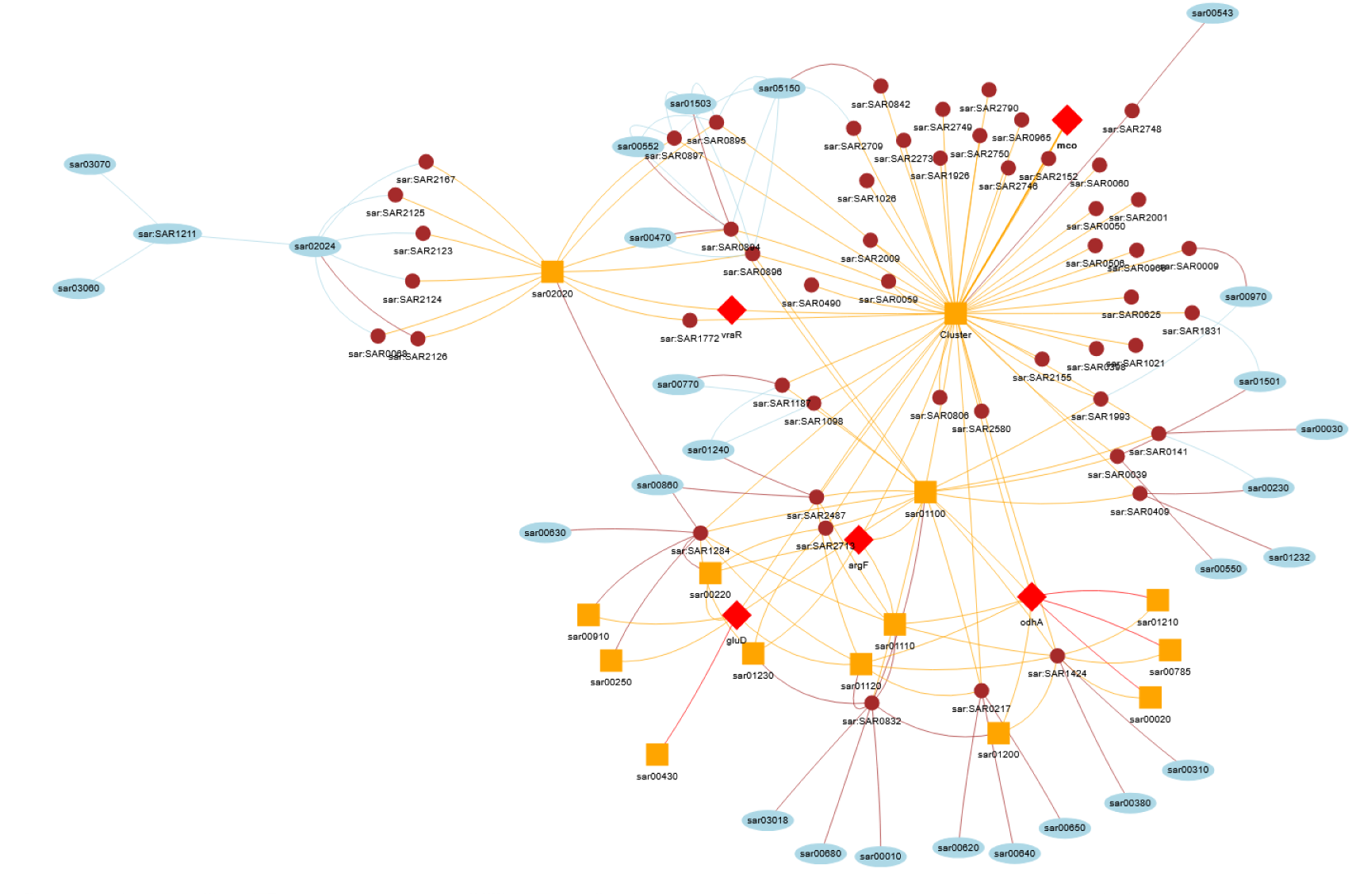
Metabolic network interconnecting genes and pathways. Nodes represent genes and metabolic pathways, while edges indicate functional associations. Genes of interest are highlighted in red with diamond shape, metabolic pathways in orange, and connected genes in brown, facilitating visual interpretation of functional structure and identification of key elements in the network.

Interactive visualization, developed with Pyvis, allowed clear distinction of node types through colors and shapes: genes of interest in red with diamond shape, pathways in orange, and connected genes in brown.

Key topological metrics were calculated for each gene in the network, including degree, betweenness centrality, and clustering coefficient. These metrics reflect the structural and functional importance of genes in the network. Then, a binary objective was defined to classify genes associated with pathways related to biofilm and virulence.

With a Random Forest classifier trained on these features, high predictive capacity was obtained, evidenced by precision, recall, and F1-score metrics close to unity.

To capture more complex contextual relationships, Node2Vec was used to generate vectorial representations (embeddings) of nodes. When training a Random Forest model with these embeddings, improved classification was obtained compared to topological metrics alone.

Finally, topological metrics were integrated with Node2Vec embeddings, enriching the vectorial representation of each gene. This combined model achieved 83% accuracy, with perfect precision for class 0 (1.00) but moderate recall (0.67). In contrast, class 1 showed perfect recall (1.00) and precision of 0.75. This asymmetry could reflect connectivity differences in the network, indicating that class 1 forms more defined communities.

Variable importance analysis showed that betweenness centrality contributed 58.4%, followed by degree with 41.6%, while clustering coefficient was not informative. This highlights the role of connectivity and strategic position of the gene in the network over local cohesion.

Genes predicted by the model as functionally relevant (class 1) were highlighted in the network using purple color, triangular shape, and increased size, facilitating their visual identification.

Confusion matrices were generated (Figure 13) for:

- Model with topological characteristics
- Model with Node2Vec embeddings
- Combined model

**Figure 13:**
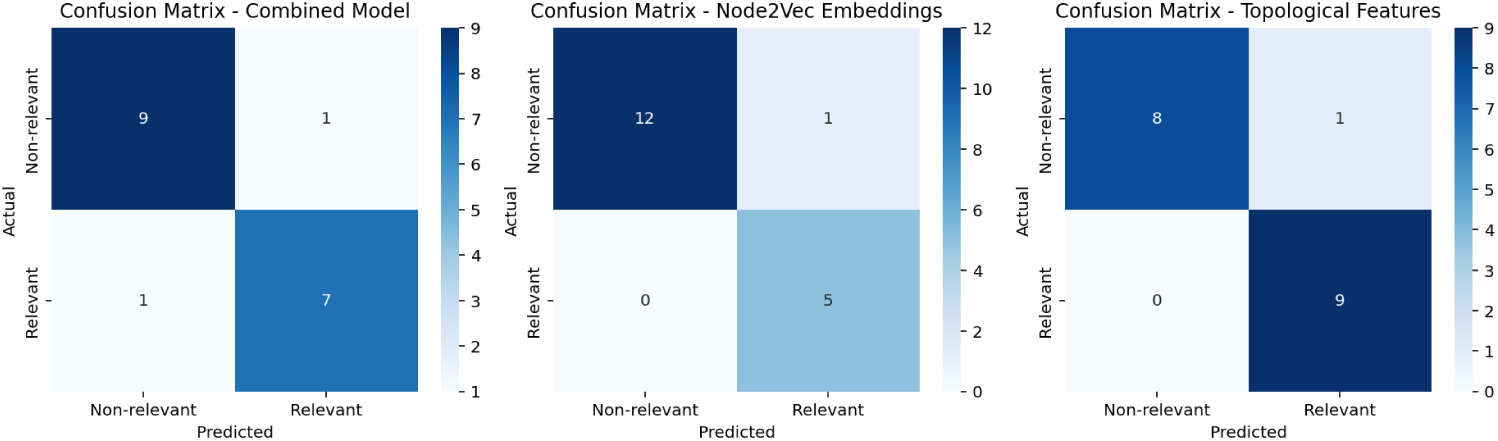
Confusion matrix comparing performance of the three models.

The topological model maximized discovery of new genes (9 TP, 0 FN), the combined model offered optimal balance (7 TP, 1 FP), and the Node2Vec model was most conservative (5 TP, 0 FN).

Metabolic pathways associated with key genes were analyzed, as can be observed in Figure 14 and in Table 3, highlighting *sar:SAR1425*, *sar:SAR0920*, and *sar:SAR1142*, linked with multiple biofilm and virulence processes. The *VraR* gene showed a notable role regulating a two-component system linked to peptidoglycan membrane.

**Figure 14:**
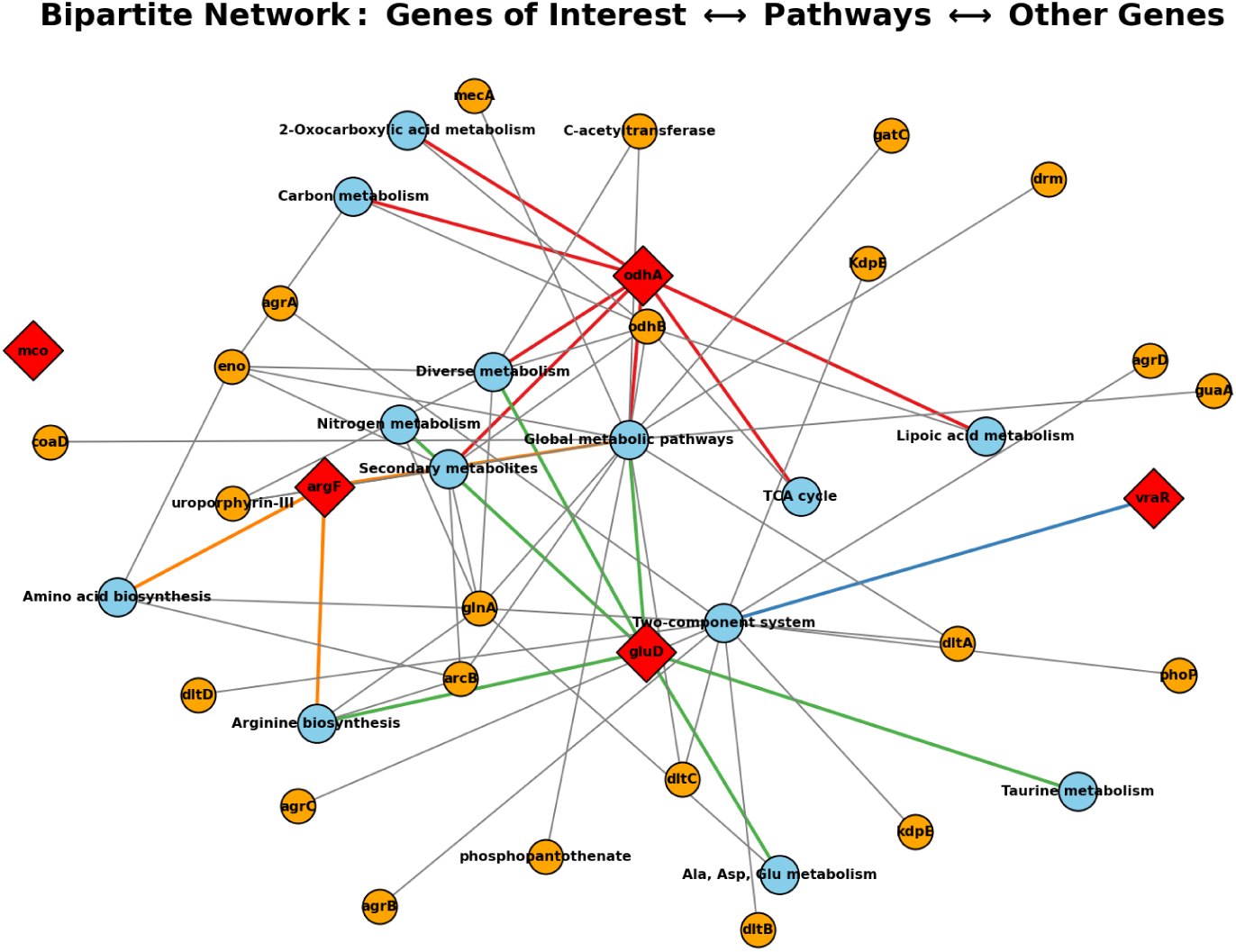
Functional network connecting genes of interest, SAR regulatory genes, and their associated KEGG pathways.

**Table 3:** Genes of interest with their associated pathways.

| Gene | Associated pathways |
| --- | --- |
| odhA | TCA, lipoic acid, 2-oxocarboxylic, carbon, secondary metabolites, diverse |
| VraR | Two-component system |
| gluD | Taurine, Ala/Asp/Glu, arginine, secondary metabolites, diverse, global |
| mco | — |
| argF | Global, amino acids, arginine, secondary metabolites |

Additionally, a dot-plot was constructed showing shared pathways between genes of interest and other connected genes, highlighting that *VraR* shares pathways with *argX*, *Vrar*, and *kdpX*, all linked with biofilm and adaptation in Figure 15.

**Figure 15:**
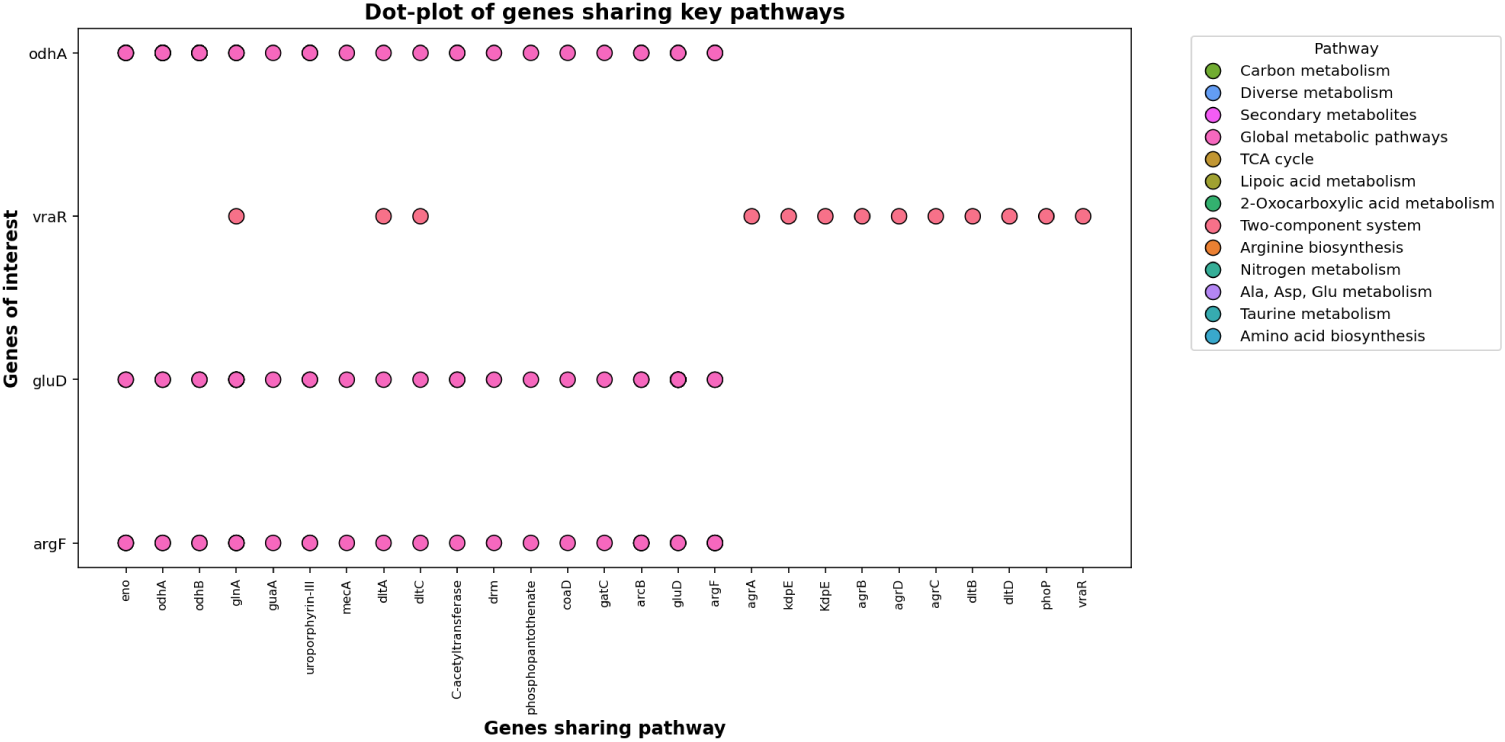
Dot-plot showing genes that share KEGG pathways with genes of interest.

## Discussion

Bacterial exposure to microgravity conditions triggers profound modifications in gene regulation, altering essential processes such as metabolism, stress response, virulence, and cellular architecture [16, 42, 18]. The differential expression analysis performed with DESeq2 revealed that genes with statistically significant regulation (p-value *<* 0.05) were concentrated exclusively in Petri dish culture conditions (BRIC-23). In contrast, both astronaut microbiota samples and spacecraft surface samples showed no changes that reached statistical significance.

This disparity in detection could be explained by differences in the intensity of environmental signals: pure culture on Petri dishes exposes bacteria to uniform osmotic stress and eliminates complex microbial interactions, creating a more direct selective pressure that facilitates the identification of specific gene responses. Conversely, in microbiota ecosystems, the presence of competing bacterial and fungal communities may mask more subtle changes through compensatory mechanisms or functional redundancy [16, 44, 42, 32, 18, 40, 24].

In the heatmaps of the most variable genes in microbiota and surface scenarios, only scarce genes with high variability were identified. Most of these genes showed underexpression, while only a minority exhibited overexpression. This pattern suggests that, although massive transcriptional change does not occur at a global level, variability arises because, under most conditions, the expression of these genes sits below their mean, presenting overexpression peaks only under very specific conditions.

In contrast, the coexpression analysis using WGCNA successfully identified robust modules specifically in Petri culture conditions, which were subjected to rigorous validation through bootstrap techniques. These modules showed notable enrichment in processes related to cell wall biosynthesis (such as the blue module, GO:0009252) and energy metabolism (exemplified by the red module, GO:0006096) [14]. However, it is striking that in microbiota and surface scenarios, correlations between module eigengenes and evaluated phenotypic traits, such as biofilm formation, did not reach statistical significance (p *>* 0.05). This finding suggests that coexpression by itself does not constitute a sufficient indicator to establish direct causal relationships in highly complex microbial environments.

This finding was reinforced with the bootstrapping technique, showing the observed differences in module stability between conditions that could be attributed to several factors. First, differential data quality between experiments can influence modular stability, where conditions with greater technical noise tend to show less reproducible patterns. Additionally, the biological variability inherent to each experimental system can contribute to these differences; for example, pure cultures like BRIC_23 typically exhibit more stable patterns than complex systems like microbiota.

Sample size also plays a crucial role, as conditions with fewer replicates may show less stable modules due to limitations in statistical power. Another determining factor is that the complexity of the biological system studied significantly influences stability, where more complex environments naturally present more variable coexpression patterns.

In the more statistical part, the *Volcano Plots* obtained for BRIC_23 reveal an interesting pattern: although global statistical significance was not achieved, certain genes such as *vraR* showed consistent trends toward underexpression in multiple experimental conditions. This phenomenon suggests that the regulation of specific genes might be modulated by common environmental factors, such as microgravity, but their precise detection requires adjustments in statistical thresholds or increases in sample size to robustly establish whether these genes experience true alterations in their expression.

Despite this limitation, the intersection analysis between genes present in modules from different experimental conditions revealed the existence of a common core of 45 genes, which includes *vraR* and *gluD*, which appeared in at least three different scenarios. These genes, characterized by showing high variability between terrestrial and space gravity conditions, could function as “master regulators” in bacterial adaptation processes, coordinating the modulation of critical metabolic pathways in a synchronized manner.

The results of functional enrichment analysis applied to the coexpression modules identified by WGCNA reveal that *S. aureus* activates gene networks specifically associated with fundamental processes for bacterial survival. Among these processes are cellular stress response (represented by genes such as *GrpE* and *DnaJ*), cell division mechanisms (with genes like *XerC* and *XerD*), energy metabolism (including *pdh* and *qoxA*), sugar transport systems (such as *crr* /*mtlA* and *mtlF*), biosynthesis of structural components (represented by *murA2* and *gcvT*), and virulence factors (including *SaeR/SaeS*, *hlgA/B/C*, *Sbi*, and *clfA*).

The ability to form biofilms on spacecraft surfaces constitutes a critical risk both for the potential obstruction of systems and for facilitating bacterial persistence [36]. The identified coexpression modules included genes directly related to cellular adhesion (such as *clfB* and *fnbA*) and extracellular matrix production (represented by *icaA*), although their differential expression only reached significance under Petri dish culture conditions. This finding suggests that, while the genetic machinery for biofilm formation is constitutively present, its effective activation depends on specific environmental signals [39, 49, 2, 14].

This transcriptional reprogramming represents an adaptive process that is triggered when bacteria face environments with stressful conditions. The magnitude of this response is evidenced by previous studies demonstrating that up to 23% of the bacterial transcriptome can experience alterations in potentially adverse environments. In particular, marked overexpression of genes encoding chaperones and stress response proteins (such as *DnaJ* and *GrpE*) is observed, probably as a compensatory mechanism to counteract the accumulation of misfolded proteins and the proteotoxic stress characteristic of the space environment [6].

On the other hand, significant underexpression of genes related to ribosomal translation and cellular biosynthesis (exemplified by *RplC* and *pdh*) is detected, suggesting a reduction in growth rates and biosynthetic activity under space conditions. Decreases have also been documented in the expression of genes involved in ionic transport and bacterial motility, possibly reflecting alterations in cellular homeostasis in the absence of gravitational gradients.

Regarding genes related to virulence and adhesion, the evidence is more variable and dependent on the specific experimental context of each study, with both increases and reductions in their expression being observed [3]. In our analysis, the identification of these genes within common modules between different experimental scenarios reinforces the hypothesis that there is a coordinated physiological adaptation, where functions related to biofilm formation, adhesion processes, and intracellular modifications remain active or experience similar regulation both in the SpaceX Inspiration4 mission and in BRIC-23.

The presence of the *murA2* gene, directly involved in peptidoglycan synthesis, indicates that modifications occur in cell wall structure when bacteria face altered conditions of mechanical pressure and gravity, as has been demonstrated in studies conducted with *Paraburkholderia phymatum* [6].

The observed underexpression of *vraR* in BRIC_23—a key component of the VraSR two-component system—coincides with previous findings documented in *S. aureus* strains subjected to osmotic stress or exposed to *β*-lactam antibiotics. This gene plays a fundamental role in regulating peptidoglycan biosynthesis and cellular response to cell wall damage. Its downregulation could represent an adaptive strategy to avoid overproduction of structural components in environments characterized by resource limitation. Simultaneously, the observed overexpression of *gluD* (glutamate dehydrogenase), *odhA* (*α*-ketoglutarate dehydrogenase subunit) and *argF* (ornithine carbamoyltransferase) suggests an increase in ATP production and essential metabolites, including glutamate and arginine, compounds critical for maintaining cell membrane integrity and polyamine synthesis under stress conditions [40].

This uncoupling between the classic stress response mediated by the VraSR system and the activation of alternative metabolic pathways could represent a strategy to optimize energy resource use under microgravity conditions, where both fluid dynamics and nutrient availability experience drastic alterations. Studies conducted with *Bacillus subtilis* have demonstrated that microgravity modifies the expression of metabolic genes, favoring fermentative pathways over aerobic respiration, a pattern consistent with the upregulation observed in *odhA* and *gluD* [18, 40].

The findings derived from the BRIC_23 experiment, which specifically analyzes the alteration of quorum sensing mechanisms, complement and reinforce our results. This experiment demonstrated that *S. aureus* under microgravity conditions experiences an increase in cell density that could explain the activation of the Agr quorum sensing system, which regulates multiple virulence factors. This phenomenon is consistent with our observation of metabolic overexpression directed toward maintaining cellular viability in restrictive environments, as evidenced by elevated levels of *gluD*, *odhA* and *argF*, fundamental genes in pathways that generate ATP and essential nitrogen compounds. Additionally, the global reduction in secreted proteins during spaceflight, attributed to high expression of proteases, fits with the need to modulate the extracellular proteome as a form of adaptive efficiency. The pattern of *vraR* underexpression observed in our analysis could also be influenced by nutrient limitation which, according to BRIC_23 suggests, is reflected in the repression of *CodY*, a global regulator sensitive to cellular metabolic state [35, 36].

These conditions favor the activation of alternative pathways, possibly more efficient in terms of biosynthesis and stress response, which is coherent with the observed uncoupling between classic cellular damage signals and the activation of fermentative or tricarboxylic acid cycle responses. The temperature of 34°C employed in BRIC_23 could have contributed to generating a gene expression profile more oriented toward persistence on surfaces, supporting the hypothesis that *S. aureus* under microgravity adopts a physiology more compatible with exogenous transmission. Together, these findings reinforce the proposal of a global reprogramming of metabolism and virulence of this bacterium under space conditions [16, 18].

The recurrent appearance of *vraR* in modules corresponding to different experimental conditions highlights its dual role in antimicrobial resistance and virulence regulation. In *S. aureus* strains with vancomycin-intermediate resistance (VISA), mutations in *vraR* are associated with alterations in membrane fluidity and anteiso fatty acid synthesis, critical components for adaptation to hostile environments. The underexpression observed in this study could have a functional effect equivalent to such mutations, reducing susceptibility to glycopeptides through modulation of cell wall biosynthesis [36, 34, 12]

Despite not having identified genes with statistically significant differential expression in microbiota samples during the space mission, it is plausible that these genes exert functional effects similar to those observed in bacterial cultures on Petri dishes, as suggested by both our findings and those reported in other studies [44].

The analysis performed by [44], which represents the most extensive dataset to date on spaceflight-associated microbiome, demonstrates significant changes at bacterial, viral, and genetic levels that show high correlations with alterations in host immune response. However, the interpretation of these results faces several important limitations: the reduced crew size (n=4), the low amounts of biomass in samples that required PCR amplification (introducing potential biases), and the inherent complexity of distinguishing between specific effects of the space environment (microgravity, radiation) and those derived from the shared environment, such as standardized diet or prolonged physical proximity between astronauts.

The *NASA Twins Study* [12] faced similar methodological difficulties due to limited sample sizes and restrictions in sampling strategies. Additionally, the detection of bacteria and other organisms such as viruses in metagenomes and metatranscriptomes presents particular technical challenges due to high diversity and limitations in available taxonomic databases [44].

Considering these limitations, although robust expressional changes were not observed in the analyzed genes, the strong correlation between microbiota modifications and immune response activation suggests that the microbial adaptations detected in bacterial cultures could be occurring *in vivo*, actively modulating host-microbiome interaction under space conditions [44]. This hypothesis requires validation through studies that incorporate larger sample sizes, improved sampling methods, and experimental models that allow isolation of specific effects of microgravity and other space factors on microbiota and host immune response.

A particularly significant finding is the discrepancy observed between the results of this study, based on 48 hours of exposure, and previous observations documented in prolonged space missions, where *S. aureus* strains developed more pronounced and stable resistances.

The early underexpression of *vraR* may represent an initial phase of metabolic adjustment preceding the activation of more permanent resistance mechanisms. This notion is supported by studies on *Pseudomonas aeruginosa* under both simulated and real microgravity conditions. The investigation [5] reported significant physiological changes during the first six days of simulated microgravity, reflecting early metabolic adaptation. Earlier studies under low shear modeled microgravity [1] and spaceflight conditions [21] demonstrated differential gene expression, regulatory shifts including sigma factor AlgU, increased cell density under nutrient-limited environments [20], and alterations in biofilm formation. Collectively, these results support a temporal adaptation model in which early metabolic reconfiguration occurs during the first 72 hours. This may be followed by the induction of resistance mechanisms, such as efflux pumps, within a week, and could ultimately culminate in chromosomal mutations over prolonged exposure periods.

The detection of genes with variable expression, such as *vraR* and *gluD*, in early stages of exposure suggests that these elements could function as “environmental sensors,” preparing the bacterium for more pronounced adaptive changes in later phases. However, the absence of longitudinal data in our study prevents confirming whether the observed trends consolidate into stable and heritable phenotypes [24].

The inherent complexity and methodological limitations of current studies on space-flight microbiome highlight the urgent need to design more controlled and larger-scale investigations to comprehensively understand microbial-immune and microbial-surface dynamics in the microgravity environment.

The absence of statistical significance in certain experimental conditions could reflect limitations in the experimental design employed. For example, in complex microbiota environments, high inter-sample variability—a consequence of multifactorial bacterial interactions—substantially reduces the statistical power of the analysis. Strategies such as implementing more standardized metatranscriptomic sequencing protocols could contribute to mitigating this problem in future research.

Additionally, the usual methodological approach that focuses on genes with *fold-change* > 2 could overlook subtle but biologically relevant regulations. In *Mycobacterium tuberculosis*, for example, changes of only 1.5-fold in the expression of regulatory genes such as *dosR* are sufficient to trigger highly important adaptive responses, such as the transition toward latent states. This suggests the need to implement more flexible and contextualized thresholds for result interpretation in contexts of moderate but sustained stress.

Therefore, it is fundamental that future studies implement more rigorous terrestrial controls, increase both cohort size and sampling frequency, and utilize integrated multi-omics approaches with controlled experimental models. These elements are essential for unraveling the specific molecular pathways that mediate bacterial responses, considering that the microbiome represents a highly complex system characterized by extraordinarily heterogeneous interactions [24].

This study demonstrates that gene expression changes in *S. aureus* under microgravity are not random, but reflect **coordinated, evolutionarily conserved adaptations** consistent across diverse spaceflight scenarios, including the ISS, human microbiota, and spacecraft surfaces. This **transcriptomic convergence** highlights the deployment of conserved response pathways that ensure survival, virulence, and persistence under extreme environmental stress. Our analysis shows that *S. aureus* prioritizes **metabolic reprogramming** over classical resistance mechanisms, exemplified by the underexpression of vraR and activation of alternative energy pathways, reflecting remarkable **regulatory and metabolic plasticity**.

These insights support practical strategies for space microbiology, including the development of molecular monitoring systems using early biomarkers (e.g., gluD, odhA), design of targeted countermeasures interfering with *anteiso* fatty acid synthesis to control membrane fluidity, and comprehensive longitudinal studies to capture the dynamics of adaptation and resistance development. Integrating transcriptomic and phenotypic analyses, such as antibiotic susceptibility, biofilm formation, and virulence assays, will be crucial for validating these strategies and ensuring astronaut health. Overall, these findings provide a foundation for next-generation microbial control approaches in space, advancing both scientific understanding and crew safety.

## Methods

### Experimental Design

#### Missions Background

The SpaceX Inspiration4 mission provided a unique opportunity to study both human microbiota and the microbial environment of spacecraft surfaces during a commercial spaceflight. For the human microbiota component, swabs were collected from ten different body regions of the four astronauts. This sampling was carried out following a rigorous temporal protocol spanning eight carefully scheduled measurement points. The pre-launch phase included three evaluations: the first conducted 92 days before takeoff (L-92), followed by a second measurement 44 days prior (L-44), and a final assessment three days before launch (L-3). During the actual space mission, two samples were obtained corresponding to flight day 1 (FD1) and flight day 2 (FD2). Subsequently, the protocol included three follow-up measurements: an immediate evaluation the day after return (R+1), a second measurement 45 days after landing (R+45), and a final evaluation 82 days post-flight (R+82). This temporal methodology generated a total of 40 biological samples that allowed for complete longitudinal analysis of the process.

Simultaneously, environmental samples were obtained from ten internal zones of the Dragon capsule at three critical moments: one pre-launch measurement (L-44) and two during flight (FD1, FD2), totaling 30 additional environmental samples. All sample collection was performed using Isohelix Buccal Mini Swabs and tubes containing DNA/RNA Shield to preserve genetic material integrity.

It is essential to highlight that all sample collection was conducted under strict informed consent and with prior approval from the ethics committee of Weill Cornell Medicine (IRB Protocol 21-05023569), thus ensuring compliance with the highest ethical standards in current biomedical research.

Total RNA extraction protocol was executed using the QIAGEN AllPrep DNA/RNA/Protein Kit, with the specific modification of omitting steps 1 and 2 of the standard protocol to optimize genetic material recovery. Cell lysis was performed through mechanical homogenization using PowerBead tubes with 0.1 mm glass beads for ten minutes, ensuring complete and uniform cell disruption.

After extraction, precise RNA quantification was performed using a Qubit 4 Fluorometer. Subsequently, processed samples were sent to HudsonAlpha facilities, where they underwent DNase I treatment to eliminate any residual DNA contamination, followed by ribosomal depletion using the NEBNext rRNA Depletion v2 (Human/Mouse/Rat) kit to enrich messenger RNA fractions.

Sequencing library construction was performed using the Ultra II Directional RNA kit with an initial amount of 10 ng RNA, employing dual indexed primers to enable sample multiplexing. Finally, all libraries were quantified and validated through Bioanalyzer and qRT-PCR to ensure quality before proceeding with sequencing using the Illumina NovaSeq 6000 platform in paired-end (PE) 150 bp format [44, 32].

The BRIC_23 mission, conducted aboard the International Space Station (ISS), constituted a complementary controlled experiment using pure cultures of S. aureus subsp. UAMS-1. Bacterial cultures were strategically distributed in four BRIC-PDFU (Biological Research in Canisters - Petri Dish Fixation Units), specialized containers designed specifically to house Petri dishes under microgravity conditions.

The experimental design included sending two units to space aboard the SpaceX-9 mission (launched July 18, 2016), while the remaining two units were kept on Earth as terrestrial controls, replicating ISS environmental conditions except for the absence of microgravity. Each experimental unit contained five Petri dishes, previously inoculated with 10^7^ cells of S. aureus UAMS-1, resulting in a total of ten spaceflight plates and ten ground control plates.

Bacterial growth was activated through controlled addition of TSY-glycerol medium, and samples were incubated for 48 hours at a temperature of 20–30°C without mechanical agitation. At the end of the incubation period, all samples were immediately frozen at −80°C to preserve their biological state. It is important to note that the ground control experiment (GC2) required repetition due to low colony forming unit (CFU) recovery in initial cultures [16].

Prior to Earth return (August 29, 2016), space samples were maintained at −32°C on the ISS. Once on Earth, they were transported to Kennedy Space Center (KSC) where they were conserved at −80°C, and subsequently processed at Ames Research Center (ARC) within NASA’s GeneLab platform. Final processing included separation of bacterial cells from plates through mechanical scraping, optical density measurement at 660 nm and estimation of total cell number for subsequent transcriptomic analysis. Sequencing was performed using the Illumina HiSeq 4000 platform in single-end (SE) 90 bp mode.

In both experiments, summarized in Table 4, a ribosomal depletion strategy was implemented to remove ribosomal RNA (rRNA) and concentrate analysis on functional transcripts, significantly improving sensitivity to detect genes expressed at low levels.

**Table 4:** Summary of the most important mission data. Own elaboration.

| Category | SpaceX Inspiration4 | BRIC_23 (ISS) |
| --- | --- | --- |
| Main objective | Study gene expression in human and environmental samples | Study gene expression in controlled *S. aureus* cultures under microgravity |
| Sample type | Human skin swabs (10 body zones, n=40) and environmental surfaces (n=30) | Petri dish cultures of *S. aureus* UAMS-1 (10 flight, 10 control) |
| Microorganism origin | Isolated from human microbiota and Dragon capsule | Laboratory-inoculated experimental strain UAMS-1 |
| Collection context | Crewed commercial mission (SpaceX Inspiration4) | Automated mission on ISS (SpaceX-9) |
| Flight conditions | Pre-flight, in-flight and post-flight (8 temporal points) | Post-incubation only, then freezing and return |
| Ground controls | Not explicit (pre/post flight comparison) | Yes, replicated ISS conditions on Earth |
| Number of samples | 70 total samples (40 human + 30 environmental) | 20 plates: 10 flight + 10 control |
| RNA extraction method | QIAGEN AllPrep with mechanical lysis (bead-beating, 0.1 mm) | Cell scraping from plates (detail not specified) |
| Pre-sequencing treatment | DNase I, NEBNext v2 ribosomal depletion, Ultra II library | Ribosomal depletion (strategy not detailed) |
| Sequencing platform | Illumina NovaSeq 6000, paired-end (2×150 bp) | Illumina HiSeq 4000, single-end (90 bp) |
| RNA quality | Quantified by Qubit, validated by Bioanalyzer and qRT-PCR | Not specified |
| Processing location | HudsonAlpha (USA) | Ames Research Center (NASA) |
| Genomic alignment | Against MRSA252 strain (high similarity with UAMS-1) | Same: aligned against MRSA252 |
| Common alignment purpose | Improve functional annotation and comparability between experiments | Same: facilitate joint analysis |
| Ribosomal depletion strategy | NEBNext rRNA Depletion v2 | Yes, without detailing method |

To date, no study has integrated the three critical spaceflight scenarios: the microbiota of the four astronauts from the Inspiration4 mission, the microbiota of surfaces inside the Dragon capsule, and *S. aureus* strains from the BRIC-23 experiment aboard the ISS (shown in Figure 16 and Table 5). Figure 16 illustrates the integrative analysis, which combines RNA-seq and metatranscriptomic data to identify common biochemical and genetic profiles across these distinct environments, with colored arrows and symbols indicating the direction of analysis and relationships between datasets. Table 5 summarizes the experimental scenarios, including *S. aureus* cultures from the BRIC-23 mission (ISS), human microbiota samples from four astronauts during the Inspiration4 mission, and environmental surface samples from the Dragon capsule. For each scenario, the number of samples and experimental conditions are specified, highlighting spaceflight versus ground controls and different sampling time points. In this work, we performed an integrative analysis of these three datasets to determine whether common biochemical or genetic profiles are shared across the different environments.

**Figure 16:**
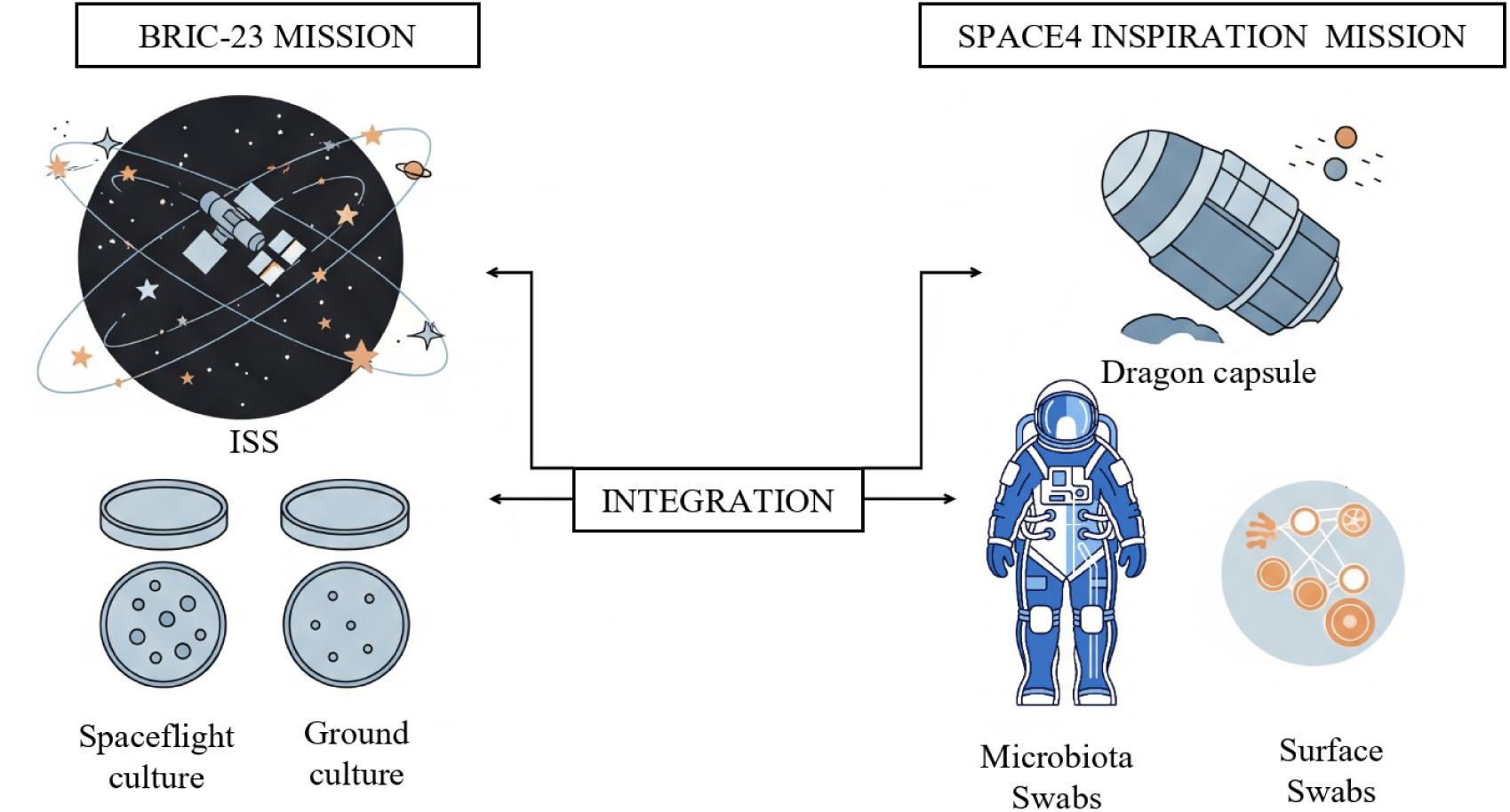
Objective of the study. This figure illustrates the integration of three critical spaceflight datasets: microbiota from the four astronauts of the Inspiration4 mission, microbiota from surfaces inside the Dragon capsule, and *S. aureus* strains from the BRIC23 experiment aboard the ISS.

**Table 5:** Experimental scenarios and sample types. This table summarizes the three spaceflight datasets integrated in the study.

| Scenario | Samples | Conditions |
| --- | --- | --- |
| ISS | 18 Petri dishes (9 space + 9 ground) | Spaceflight vs. Ground (frozen) |
| Inspiration4: Microbiota | 10 body sites × 4 astronauts (n=40) | Preflight, Inflight2, Inflight3, Postflight |
| Inspiration4: Surfaces | 10 Dragon capsule zones × 3 conditions (n=30) | Preflight, Inflight2, Inflight3 |

#### Bioinformatics Pipeline Development

Bioinformatics processing of samples was performed through custom *pipelines* developed in Bash, designed specifically to handle the inherent complexity of data generated in this study. The diversity of experimental conditions, including environmental microbiota, treated surfaces and BRIC-23 space culture samples, required a flexible computational strategy that allowed parallel processing of independent projects, as well as integration of diverse cleaning, filtering and quantification methods.

The *pipelines* were conceptualized with modular architecture that facilitated code block reuse between different experimental scenarios (microbiota/surface and BRIC_23). This modularity not only guaranteed methodological consistency throughout the study, but also provided technical adaptability shown in Figure 17

**Figure 17:**
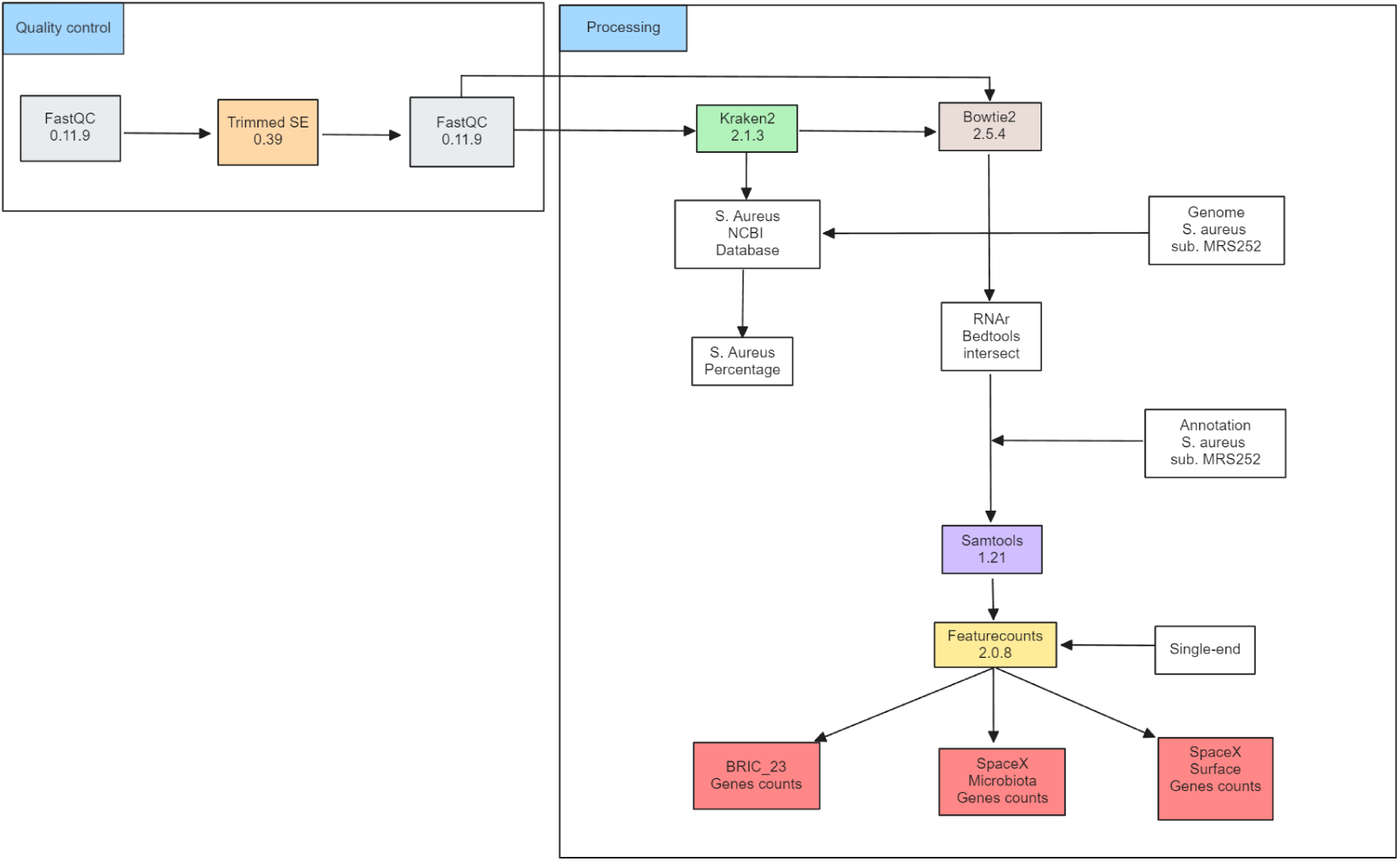
Diagram of the curation process with quality control and data processing in bash on Linux, using an automated workflow.

Development of these custom scripts provided precise control over parameters of each tool used, including quality thresholds, alignment sensitivity and confidence levels in taxonomic classification. This flexibility allowed iterative parameter adjustment according to specific requirements of each dataset, thus optimizing quality and reproducibility of results.

Initial quality control of raw sequences was performed using FastQC v0.11.9, a tool that evaluates sequencing read quality through a series of standardized metrics [17]. FastQC identifies common problems in sequencing data, including residual adapters, low-quality sequences, loss of base randomness, or systematic biases introduced during the sequencing process.

In the developed pipeline, this analysis was systematically executed for all raw sequence samples, generating detailed reports that were organized into specific folders ($FASTQC_DIR/original/$SAMPLE_NAME). This organization facilitated systematic visual review and early identification of potential technical problems before proceeding with any additional bioinformatic processing.

Subsequently, Trimmomatic v0.39 was implemented for read trimming, constituting a critical step to improve sequence quality before alignment and gene quantification [46]. Trimmomatic was configured to automatically remove ILLUMINA and NEXTERA adapters using predefined adapter files in the software.

The specific parameters used were:

- ILLUMINACLIP:“$ILLUMINA_ADAPTERS“:2:30:10:8:true: Illumina adapter trimming with configuration allowing a maximum of 2 mismatches, establishing a minimum quality threshold of 30, defining a minimum length of 10 nucleotides for valid trimming, and specifying 8 bases to trim from the ends when the adapter is recognized.
- ILLUMINACLIP:“$NEXTERA_ADAPTERS“:2:30:10:8:true: Nextera adapter trimming using identical parameters to those used for Illumina adapters.
- SLIDINGWINDOW:4:20: Implementation of a sliding window algorithm that evaluates groups of 4 consecutive bases, removing segments where average quality is below 20. This approach is particularly effective for eliminating low-quality regions that could compromise subsequent analyses.
- MINLEN:50: Removal of reads that, after the trimming process, show a length below 50 base pairs, thus avoiding retention of fragments too short that could generate ambiguous alignments.

This processing was fundamental for reducing errors in subsequent alignment and quantification stages, considering that reads with residual adapters or low quality can generate mapping errors, increase false positives in gene quantification, and generally significantly compromise the accuracy of final results.

The processing was adapted to the specific technical characteristics of each dataset: SpaceX samples, characterized as paired-end (PE), were processed using Trimmomatic’s PE option, allowing simultaneous trimming of both files (R1 and R2) while preserving only high-quality paired reads. In contrast, BRIC_23 samples, being single-end (SE), were processed through the SE option, trimming exclusively one sequence file per sample. Once the trimming process was completed, a second quality analysis was performed using FastQC on the processed reads ($FASTQC_DIR/trimmed/$SAMPLE_NAME). This validation analysis confirmed substantial improvement in sequence quality, demonstrating the effectiveness of the preprocessing procedure. The generated reports showed that reads had significantly fewer residual adapters and better overall quality scores, ensuring that sequences were prepared for subsequent alignment and quantification analyses.

In the processing phase, Kraken2 v2.1.3 was employed to perform taxonomic classification of preprocessed reads, using a specialized *S. aureus* database (taxID 1280) [13]. This analysis allowed precise quantification of the percentage of reads assigned to this specific species in each analyzed sample.

Kraken2 was configured to execute taxonomic classification based on fixed-length *k-mers* of 35 base pairs, using a predefined database that includes reference genome sequences from a wide diversity of microbial species, but was limited to S.aureus. The specific parameters used included:

- –confidence 0.2: to establish the minimum confidence threshold required for taxonomic assignment.
- –threads: to optimize performance through analysis parallelization.

It’s important to note that systematic discrepancies were observed between results obtained through Kraken2 and those generated by Bowtie2 v2.5.4 [4]. This divergence is explained by fundamental differences in the methodologies employed: Kraken2 uses an approach based on short *k-mers* and general taxonomic databases, while Bowtie2 implements precise and specific alignment against the *MRSA252* reference genome.

For genomic alignment, preprocessed reads were aligned against the complete *S. aureus MRSA252* genome using a pre-built index with Bowtie2. Specific alignment parameters included:

- -L 20: adjustment of minimum length of alignment fragments (seed length).
- -k 1: limitation of the number of alignments reported per read to one, prioritizing the highest quality alignment.
- -p $THREADS: enabling the use of multiple processing threads to optimize computational performance.

This configuration allowed highly specific mapping of reads against the *MRSA252* reference genome. The observed discrepancies between Kraken2 and Bowtie2 are primarily attributed to differences in handling reads with imperfect alignments or genomic variants not represented in the Kraken2 taxonomic database.

Aligned sequences were processed using the Samtools suite of tools for conversion and manipulation of alignment files. SAM files generated by Bowtie2 were converted to BAM format using samtools view, followed by sorting through samtools sort to optimize access and subsequent processing.

Subsequently, Bedtools intersect was employed to intersect aligned reads with gene annotations of the reference genome. This process allowed overlapping reads with genomic regions of interest, defined by annotations in GFF format containing detailed information about exact gene positions and functional characteristics. As a result, gene-level annotated BAM files were generated that included both reads aligned with coding regions and those corresponding to other functional genome regions, thus facilitating subsequent gene expression quantification.

Gene expression quantification was performed shown in a diagram at the Figure 17. Starting using FeatureCounts v2.0.8, configured to use annotated BAM files as input and running in single-end mode, even for originally paired-end samples [38]. This methodological decision was based on preliminary analyses that revealed specific mapping patterns according to sample type.

After initial preprocessing and removal of adapters and ribosomal RNA through ribodepletion with SortMeRNA, samtools flagstat v1.18 was employed to quantify filtration efficiency and characterize in detail the percentage of reads aligned to the *S. aureus* MRSA252 genome.

In microbiota samples (e.g., GLDS-564, Astronaut 1), analysis of the non-ribosomal reads file (_non_rRNA.bam) revealed a total of 46,890,597 processed reads. Of these, only 261,897 reads (0.56%) aligned perfectly to the *MRSA252* genome. In contrast, the ribosomal reads file (_rRNA.bam) showed 4,428,547 mapped reads (100%), with 87.15% correctly paired.

This mapping pattern reflects that, in microbiota samples, the vast majority of non-ribosomal RNA comes from numerous bacterial species whose sequences show no significant homology with the *MRSA252* genome. Therefore, the low mapping percentage is not due to deficiencies in the trimming process or pair fragmentation, but to the inherently diverse taxonomic composition of the microbiota.

In contrast, BRIC_23 samples (GLDS-145) showed a radically different mapping pattern. The *_non_rRNA.bam file contained 10,686,642 reads, of which 10,595,682 (99.15%) aligned perfectly to the *MRSA252* genome. Only 10,717 reads (100%) were found in the *_rRNA.bam file, confirming ribodepletion efficiency and culture purity.

This confirms that, in a pure culture system, most non-ribosomal RNA effectively comes from the reference strain, thus validating both culture quality and bioinformatic processing.

Given that these mapping differences reflect the natural taxonomic diversity of microbiota versus monoculture purity, the methodological decision was adopted to maintain quantification in single-end mode for all samples (both PE and SE). This strategy allowed standardization of the analysis and avoidance of biases derived from technical unpairings that could introduce artificial variability in comparisons between experimental conditions. Post-alignment quality controls were fundamental for validating rRNA filtration efficiency and providing the empirical basis for subsequent methodological decisions. These analyses ensured implementation of a reproducible and robust pipeline capable of handling the inherent complexity of different biological sample types.

Finally, the generated gene counts were systematically organized into three independent matrices: *BRIC_23*, *Microbiota* (per astronaut), and *Surface*. These data were integrated into a structured database to ensure complete analysis traceability and facilitate result reproducibility.

To guarantee the quality and integrity of analyzed data, a quality control protocol was implemented through structured MySQL database queries. The established inclusion criteria contemplated exclusively those samples that demonstrated 100% genomic assignment through the Bowtie2 algorithm, as well as gene annotation efficiency superior to 90%. Samples that did not satisfy these methodological standards were excluded from the dataset to preserve statistical validity and ensure robustness of subsequent transcriptomic analyses. The MySQL scripts used for implementing these quality filters are documented and available in the project repository.

This filtering strategy resulted in a reduction of experimental group sizes, optimizing data quality at the cost of statistical power. For example, the group corresponding to Astronaut 1 was reduced from *n* = 38 to *n* = 33 samples, while the Surface samples group decreased from *n* = 30 to *n* = 26 samples. Additionally, sampling points with equivalent temporal duration (∼48 hours) were selected between microbiota and BRIC_23 samples, ensuring coherent and methodologically valid temporal comparison.

#### Modules and Biological Function

Differential expression analysis constituted one of the fundamental pillars of this research. To address the complexity of data obtained in each experimental scenario (BRIC_23, microbiota, and surface), an analytical strategy was implemented that allowed independent examination of each condition among the scenarios, subsequently facilitating systematic comparison of results through intersection of identified genes.

Statistical processing of data was carried out using the R programming language, specifically utilizing the DESeq2 package, a tool that allows modeling of inherent variance in gene expression count data. This package is especially useful for RNA-seq data analysis, as it allows appropriate handling of the non-normal distribution typical of this type of data. After completing the normalization process, a variance stabilizing transformation (VST) was applied, a technique that is essential for reducing the effect of overdispersion characteristic of massive sequencing data and, consequently, improving the quality of subsequent visualizations.

During the preprocessing phase, the parameter sfType = “poscounts” was implemented within the DESeq2 framework, a critical methodological decision for appropriate handling of zero values present in microbiota and surface datasets. This problem represents one of the most frequent challenges in analyzing datasets characterized by low sequencing depth or high degree of data dispersion. Implementation of this parameter allowed obtaining more robust and reliable estimates. Furthermore, a comparative evaluation was performed between normalization factors obtained through the standard DESeq2 method and those based on the *poscounts* approach, observing that the latter provided considerably more stable estimates under the analyzed experimental conditions.

For exploration of underlying data structure, dimensionality reduction techniques were implemented, specifically PCA (Principal Component Analysis) and UMAP (Uniform Manifold Approximation and Projection). These analyses allowed visualization of natural clustering formation among samples according to their experimental conditions, revealing differentiated patterns in each evaluated scenario. Construction of heatmaps complemented these analyses, providing detailed visual representation of gene expression variability patterns between different experimental conditions. It’s important to note that these heatmaps focused on evaluating expression variability patterns between conditions, rather than absolute expression of individual genes, because expression levels in microbiota and surface samples were insufficient for this specific purpose.

Implementation of the WGCNA (*Weighted Gene Co-expression Network Analysis*) method represented an additional crucial component of the analysis, oriented toward identification of gene co-expression modules and detection of *hub-genes*, that is, genes that present a high degree of connectivity within each functional module. This methodological approach allows identification of gene groups that exhibit coordinated expression patterns, which frequently reflects their participation in common biological processes or shared regulatory networks.

WGCNA analysis used the VST-normalized expression matrix as input, from which a co-expression network based on topological similarity (TOM - *Topological Overlap Matrix*) was constructed. The resulting dendrogram from hierarchical analysis of genes facilitated identification of functional modules, each encoded through specific chromatic assignment. Subsequently, each module was correlated with corresponding experimental conditions (e.g., *spaceflight* group versus *ground control in BRIC_23*) through calculation of eigengene correlations, which allowed identification of treatment-dependent co-expression patterns.

Analysis results were exported at two complementary information levels: (1) eigengenes of each module, which provide a statistical summary of joint expression of each functional gene group, and (2) detailed gene composition per module, thus facilitating more specific subsequent analyses and prioritization of gene candidates for experimental validation.

The evaluation of robustness and stability of gene co-expression modules identified through WGCNA constitutes a critical aspect for validating the reliability of obtained results. To address this need, a bootstrapping procedure based on random resampling was implemented, a widely recognized statistical technique for evaluating stability of complex data structures [15, 45]. This methodological approach allows evaluating whether genes tend to cluster consistently in the same functional modules across different data subsets, thus providing robust internal validation of the inferred modular structure.

The bootstrapping procedure was implemented through five independent iterations of random sampling on the gene expression dataset datExpr_inflight. In each iteration, 80% of available samples were randomly selected, maintaining the necessary statistical representativeness for analysis. However, for specific analysis of the “Surface” dataset, due to its considerably smaller sample size, it was necessary to implement a methodological adaptation, using a lower sampling proportion adjusted to the specific characteristics of this dataset, with the objective of preserving statistical representativeness and avoiding loss of biologically relevant information.

In each iteration of the bootstrapping procedure, adjacency matrices and TOM similarity were independently recalculated, which constitute the foundations of gene coexpression analysis. From these matrices, a hierarchical dendrogram was constructed reflecting similarity relationships between genes. Functional modules were defined through application of the cutreeDynamic algorithm, implementing parameters specifically adjusted to optimize detection of smaller modules (deepSplit = 2, minClusterSize = 5), thus ensuring greater sensitivity in identifying biologically relevant modular structures.

Colors assigned to modules obtained in each iteration were systematically stored and used to construct a co-clustering matrix. In this matrix, each element (*i, j*) represents the frequency with which a specific gene pair was assigned to the same functional module across different iterations of the bootstrapping procedure. This quantification provides an objective measure of gene clustering consistency.

The co-clustering matrix was normalized by dividing each element by the total number of iterations performed, thus obtaining a probabilistic measure of co-assignment ranging between 0 and 1. Visualization of this information was performed through a hierarchical heatmap, facilitating visual identification of stable gene blocks that appear consistently clustered. This graphical representation allows clearly discerning which modules present greater structural stability and, consequently, greater reliability from a statistical perspective.

At the same time, appearance frequencies per module were calculated for each individual gene, which allowed extraction of a gene subset exhibiting modular stability. For this purpose, a stability threshold was established that was initially set at 0.8, representing a rigorous statistical criterion. However, this threshold was subsequently adjusted to 0.5 specifically for the “Surface” dataset, due to high variability observed in this dataset, where smaller modules required more flexible criteria to retain biologically relevant signals without compromising statistical robustness of the analysis.

In the specific case of microbiota 1, only one module met the stability criteria defined through the bootstrapping procedure. For this particular module, a detailed functional enrichment analysis was proceeded with, using the KEGG (*Kyoto Encyclopedia of Genes and Genomes*) database in combination with annotation information contained in the GFF file corresponding to coding sequences (*CDS*).

The functional enrichment process was initiated with implementation of a script developed in R, specifically designed to map gene identifiers to their corresponding references in refseq_protein format (wp_XXX). This step was fundamental for establishing the necessary connection with the UniProt database, allowing subsequent transformation of these references to their corresponding UniProt accessions. This identifier conversion constitutes an essential step in the analysis pipeline, as it facilitates inference of specific biological functions for each gene through access to standardized functional annotation databases.

#### Conserved Gene Analysis Across Experimental Conditions in Synthetic Biology

A particularly relevant aspect of this study consisted of analyzing whether metabolic pathways associated with genes of interest are functionally related to biofilm formation processes. For this purpose, a systematic comparison was implemented between metabolic pathways identified in our analyses and a set of pathways previously characterized and documented for their direct implication in this specific biological process. Comparative analysis revealed which genes of interest were, effectively, associated with metabolic pathways linked to biofilm formation. Although not all studied genes showed this association with known biofilm-related pathways, those that did exhibit this connection could play a crucial role in regulation and modulation of biofilm formation processes in studied organisms shown in Figure 18.

**Figure 18:**
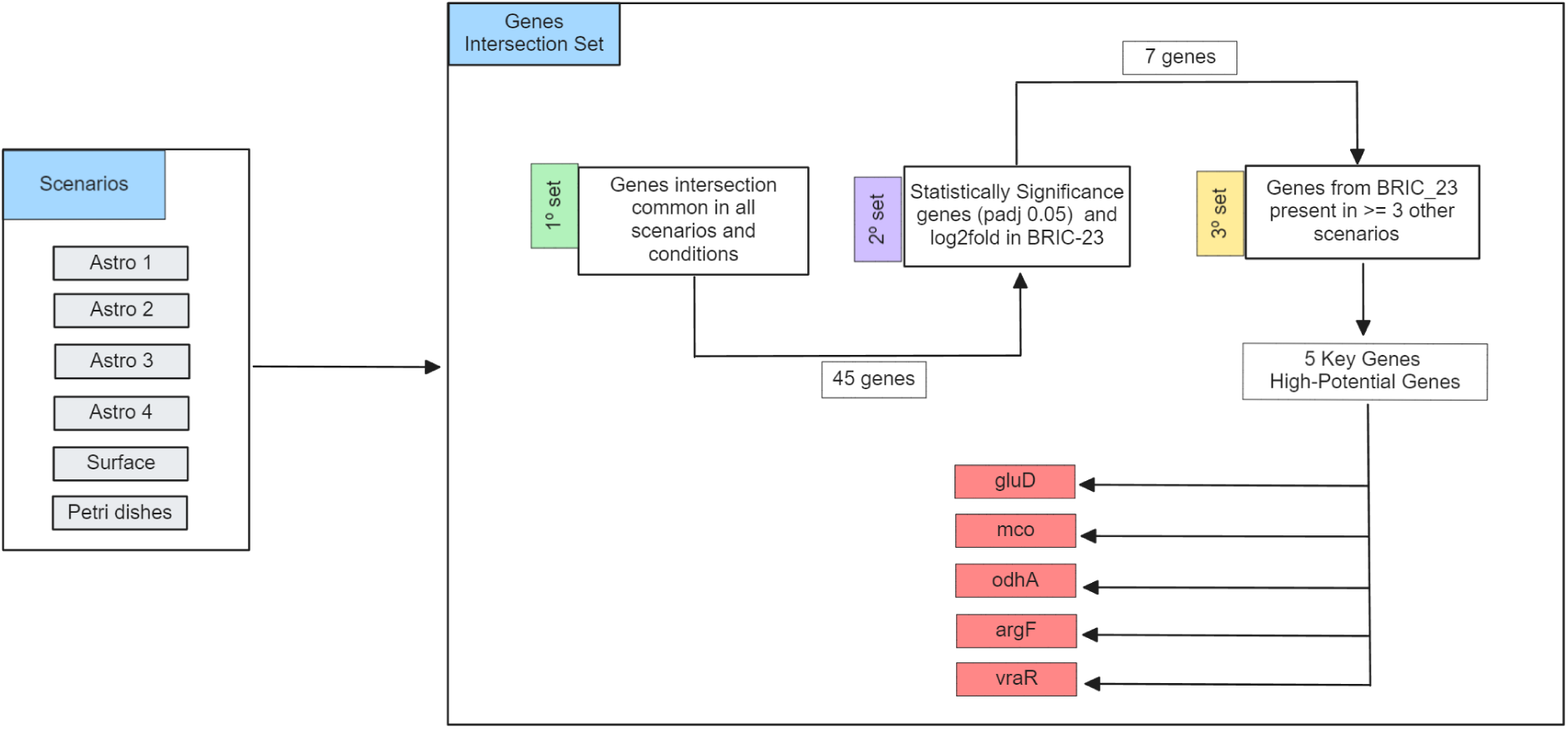
Workflow diagram of the computational pipeline used to identify genes of interest. The process includes loading CSV files for all experimental scenarios, structuring gene-module associations, identifying common genes across conditions, defining selection sets based on statistical significance and functional robustness, performing intersections, and applying additional filters to obtain the final set of robust and biologically relevant genes.

Identification of shared genes between diverse experimental conditions constitutes a fundamental step for understanding conserved biological mechanisms that could be involved in adaptive response of studied organisms. To address this objective, a computational pipeline was developed in Python, leveraging the capabilities of the pandas library for efficient handling of complex data structures.

The analysis was systematically applied to six distinct biological scenarios, including four conditions corresponding to specific microbiotas (designated as microbiota1 through microbiota4) and two additional conditions representing differentiated culture environments: Petri dish culture and spacecraft surface cultures. This diversity of experimental conditions allowed evaluation of gene expression pattern robustness across different environmental contexts.

Information from each experimental scenario was organized into structured CSV files, containing the association between gene identifiers and functional modules in two defined columns. To facilitate computational processing of these data, a function called cargar_genes_y_modulos() was implemented, specifically designed to optimize information loading and structuring. This function, when receiving the path to a specific CSV file as an argument, returns a Python dictionary where keys correspond to unique module identifiers and associated values are lists containing genes belonging to each functional module. This data structuring significantly facilitated subsequent comparison operations between different experimental conditions.

Once data loading was completed for all experimental scenarios, implementation of the common gene identification algorithm proceeded. For this task, a specialized function called encontrar_genes_comunes_todos() was defined, which accepts multiple gene dictionaries organized by modules as arguments and executes an iterative intersection process between all present gene sets. This algorithmic procedure guarantees that only those genes that appear consistently in all analyzed experimental conditions are conserved, thus providing a robust measure of transcriptional conservation.

The result of this intersection process was stored in a Python set data structure, which allowed multiple analytical operations:

- Precise quantification of the total number of common genes identified between all conditions.
- Visualization of specific identifiers of said genes.
- Export of results in CSV format for subsequent analysis or integration into other analytical platforms.

With the purpose of identifying those genes that present the greatest biological potential and demonstrate consistent functional robustness, an intersection analysis was carried out between three key gene sets, each selected through complementary but rigorous criteria shown in Figure 18.

- **First set:** The analysis focused on the list of 45 common genes identified in all experimental conditions.
- **Second set:** Genes previously identified as statistically significant, that is, those presenting biologically relevant log_2_*FoldChange* values and an adjusted p-value (*padj*) below 0.05.
- **Third set:** Genes belonging to the *Petri* condition that, additionally, demonstrated consistent presence in at least two other experimental conditions in other scenarios, thus fulfilling a rigorous criterion of inter-conditional robustness (presence in ≥ 3 experimental conditions).

Intersection analysis among sets allowed identification of genes that simultaneously satisfy the selection criteria. The resulting subset represents genes that are both statistically significant and functionally robust across diverse experimental conditions. This information was documented and exported in the genes_comunes_interseccion.csv file to facilitate its accessibility and traceability.

The analysis was executed on WGCNA modules subjected to bootstrapping with an 80% proportion (except for “Surface” where 50% was applied), systematically organized in specific directories per experimental scenario: Microbiota 1-4, Surface and Petri. Gene information was organized in a dictionary-type data structure that meticulously recorded the experimental conditions in which each specific gene appeared.

Once all information was collected and organized, a selection filter was implemented that identified those genes that simultaneously met two criteria:

- Confirmed presence in the “Petri” condition
- Additional appearance in at least two other experimental conditions (≥ 3 conditions in total)

This threshold of three conditions was established to guarantee a minimum criterion of transversal biological robustness, prioritizing identification of potentially relevant genes in multiple simulated experimental environments.

#### Networks and Metabolic Pathways

In this phase of analysis, a selection strategy directed toward genes that have been previously associated with critical pathogenic characteristics in *S. aureus* was implemented, specifically those related to virulence, antibiotic resistance, and biofilm formation. This association not only contributes to deeper understanding of each gene’s role in cellular metabolism, but also suggests possible critical metabolic pathways for selected genes, opening new avenues for future research and providing specific directions for experimental validation studies.

This selection was based on an exhaustive review of scientific literature that consistently documents the implication of these genes in pathogenic properties of the bacterium. Among selected genes were included in Appendix C all recognized for their documented participation in virulence mechanisms, antimicrobial resistance, and biofilm formation processes [19].

A systematic query to the KEGG API was implemented to obtain detailed information about metabolic pathways associated with previously selected genes. This query provided comprehensive information about metabolic pathways in which each of these genes participates, allowing establishment of direct and well-founded associations between specific genes and biological processes of interest, including virulence, antibiotic resistance, and biofilm formation.

Each gene was individually consulted in the KEGG database through its unique identification code, and associated metabolic pathways were systematically extracted from each query result. Obtained data were processed and stored in a structured manner to facilitate subsequent analysis and guarantee integrity of collected information.

From information about metabolic pathways associated with each gene, a binary matrix was constructed that precisely reflects presence or absence of each metabolic pathway in selected genes. In this matrix, rows represent individual genes and columns correspond to specific metabolic pathways. A value of “1” in the matrix indicates that the gene is functionally associated with the corresponding metabolic pathway, while a value of “0” signals absence of such functional association.

Construction of this binary matrix facilitated a simplified but precise representation of complex relationships between genes and metabolic pathways, significantly optimizing subsequent data analysis. During this process, special attention was paid to metabolic pathways directly associated with virulence processes, antibiotic resistance, and biofilm formation, given their relevance to research objectives.

To identify common functional patterns in genes associated with specific metabolic pathways, a clustering analysis was implemented using the K-means algorithm. The main objective of this analysis consisted of grouping genes based on their association profiles with metabolic pathways, thus allowing identification of genes that share similar functional characteristics and potentially related biological roles.

The optimal number of clusters was determined through an empirical process that evaluated different configurations, and an optimized implementation of the K-means algorithm applied on the binary matrix of genes and metabolic pathways was used. Clustering results allowed identification of gene groups with similar profiles in terms of their participation in specific metabolic pathways, suggesting similarities in their biological and functional roles, providing valuable information for biological interpretation of results.

With the objective of determining which genes possess significant impact on virulence processes, antibiotic resistance, and biofilm formation, a classification model based on *Random Forest* was implemented. This model allowed prediction and quantification of each gene’s relevance for mentioned biological properties, using as features the associations between genes and identified metabolic pathways.

Genes were labeled according to their documented relationship with processes of interest: those genes linked with virulence, antibiotic resistance, and biofilm formation received a positive label (1), while remaining genes obtained a negative label (0). Subsequently, the dataset was strategically divided into a training set and a test set, maintaining representativeness of both classes. The *Random Forest* model was trained using features derived from the binary matrix of genes and metabolic pathways.

Model performance evaluation was performed through key statistical metrics, including precision, sensitivity, and specificity, providing a comprehensive evaluation of the model’s predictive capacity. Obtained results allowed identification of key genes that play an important role in pathogenic mechanisms of *S. aureus*, significantly contributing to understanding of molecular biology of this pathogen.

Obtained results were subjected to a rigorous validation process through implementation of various complementary statistical techniques. Result presentation was performed through specialized visualizations, including scatter plots and confusion matrices, which provided clear and accessible representation of findings. This methodological approach not only allowed confirmation of reliability of implemented clustering and classification models, but also facilitated more intuitive and accessible interpretation of results for the scientific community.

In this phase of analysis, a specialized predictive model based on feature importance evaluation was used to quantify the impact of individual genes on specific metabolic pathways. The importance of each feature, represented by genes, was determined through implementation of the trained model, which assigned each gene a quantitative score reflecting its relevance in predicting specific metabolic pathways.

To obtain these importance scores, the feature_importances_ attribute of the trained Random Forest model was used. This value provides a quantitative measure of each gene’s contribution to the model’s predictive power. Results revealed which specific genes are most relevant for metabolic pathways under analysis, offering a clear and quantified perspective of genes that should be prioritized in subsequent studies and experimental validations.

The next methodological step consisted of associating genes of interest with specific metabolic pathways in which they are functionally involved. This process allows visualization and understanding of how each gene influences metabolic behavior of the studied cell or system. Each gene possesses a specific set of associated metabolic pathways, and this relationship was identified through consultation of a structured dataset where genes were annotated with metabolic pathways in which they participate. A set of genes of interest was selected and systematically mapped to metabolic pathways they share, allowing obtaining an exhaustive list of metabolic pathways associated with each specific gene.

To visualize relationships between genes, metabolic pathways, and other associated genes, a bipartite graph was constructed. This graph shows two types of nodes: genes and metabolic pathways. Edges between these nodes indicate that a gene is associated with a metabolic pathway, facilitating understanding of interactions between system components. Graph visualization revealed a structure of relationships between key genes, metabolic pathways, and other implicated genes, providing clear representation of how genes of interest are distributed across different metabolic pathways and how these interconnect with other genes outside the main interest set. This type of graphical representation is valuable for identifying association patterns between genes and metabolic pathways, and for signaling possible areas of interest for future research on metabolism and biofilm formation.

#### Data Availability

To ensure consistency and robustness of bioinformatics analyses, the following genomic reference files from *Staphylococcus aureus* MRSA252 strain were used:

- *S. aureus* MRSA252 GFF file: https://ftp.ncbi.nlm.nih.gov/genomes/all/ GCF/000/011/505/GCF_000011505.1_ASM1150v1/
- *S. aureus* MRSA252 genome in .fna format: https://ftp.ncbi.nlm.nih.gov/ genomes/all/GCF/000/011/505/GCF_000011505.1_ASM1150v1/

Although BRIC_23 originally used the UAMS-1 strain, we made the methodological decision to align all reads against the MRSA252 reference genome. MRSA252 was selected because it has a fully assembled genome, more complete functional annotation, and shares high genomic similarity (*>*96%) with UAMS-1, ensuring precise alignment and reliable functional annotation for comparative analyses [16, 28].

To maintain experimental consistency, only SpaceX mission samples replicating BRIC_23 temporal conditions were selected, specifically those with similar exposure time (48 hours,

“Spaceflight vs. Ground” condition). For surface samples, temporal points L-44, FD2, and FD3 were used, while for microbiota samples points L-3, FD2, FD3, and R+1 were employed. This strategy focused on the acute short-term transcriptomic response of *S. aureus*, maintaining temporal coherence with BRIC_23 and enabling methodologically sound comparisons between experiments.

#### Data repositories

- SpaceX Inspiration4 mission raw data: https://osdr.nasa.gov/bio/repo/data/ studies/OSD-572
- BRIC-23 mission raw data: https://osdr.nasa.gov/bio/repo/data/studies/ OSD-145

**Scripts repository:** https://github.com/xostc/TFM_NASA

Computational pipelines, analysis scripts, and files generated during this study are available in the indicated repository. It includes both raw data and metadata required for complete analysis reproduction, along with detailed technical documentation specifying dependencies, software versions, and an integral methodological guide for reproducible execution of all analytical procedures implemented in this research.

## Appendix A Variable Gene Heatmaps

The following presents the heatmaps corresponding to the genes with the highest variability in Microbiota scenarios 1 to 4, as well as in Surface. These figures are included in the appendices, given that they do not provide essential information for the main analysis. In general, these scenarios did not show marked transcriptional variability, unlike what was observed in BRIC_23.

In particular, the microbiota and surface datasets revealed only a reduced number of genes with high variability. Most of these genes presented levels of underexpression (represented in blue), while only a few exhibited overexpression peaks (in red). This pattern indicates that, while extensive transcriptional changes do not occur at a global level, the detected variability is due to generally low expression under most conditions, with specific overexpressions under particular situations.

**Figure 19:**
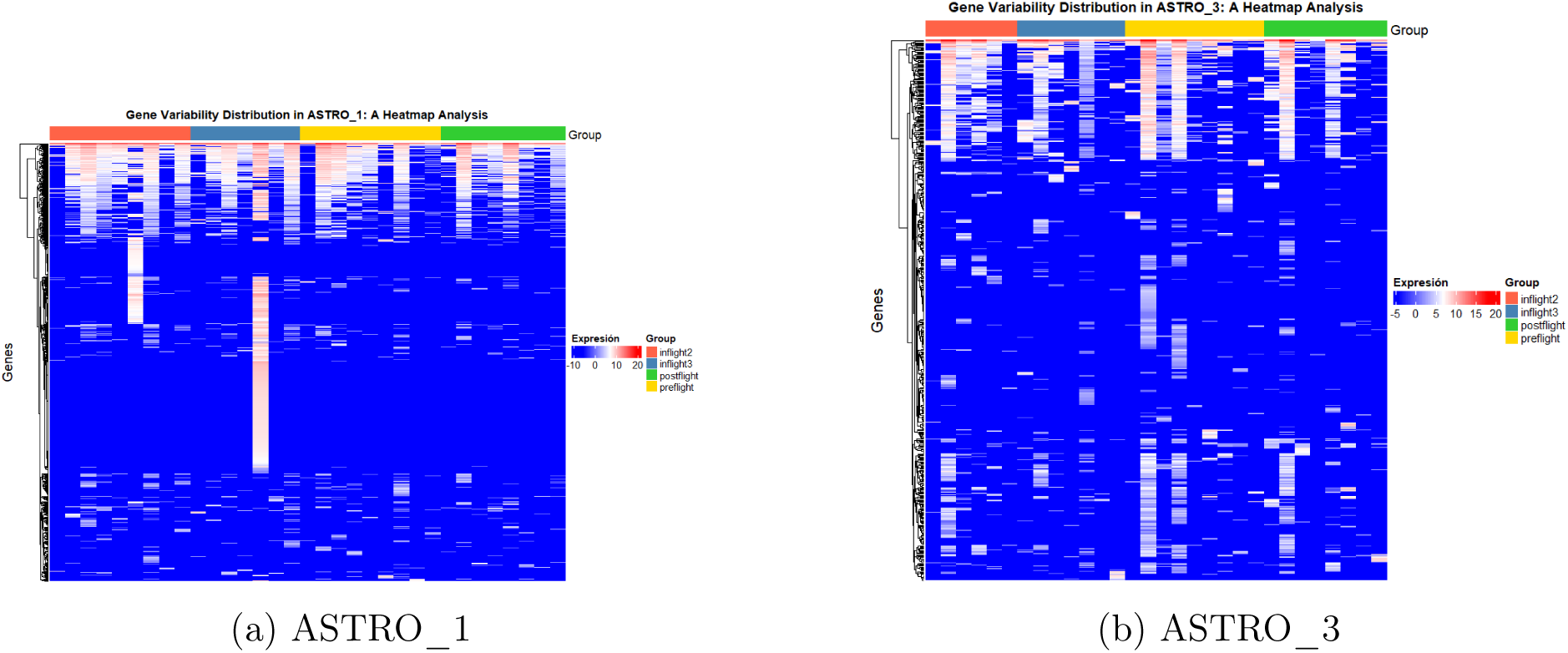
Heatmaps of most variable genes in scenarios ASTRO_1 and ASTRO_3.

**Figure 20:**
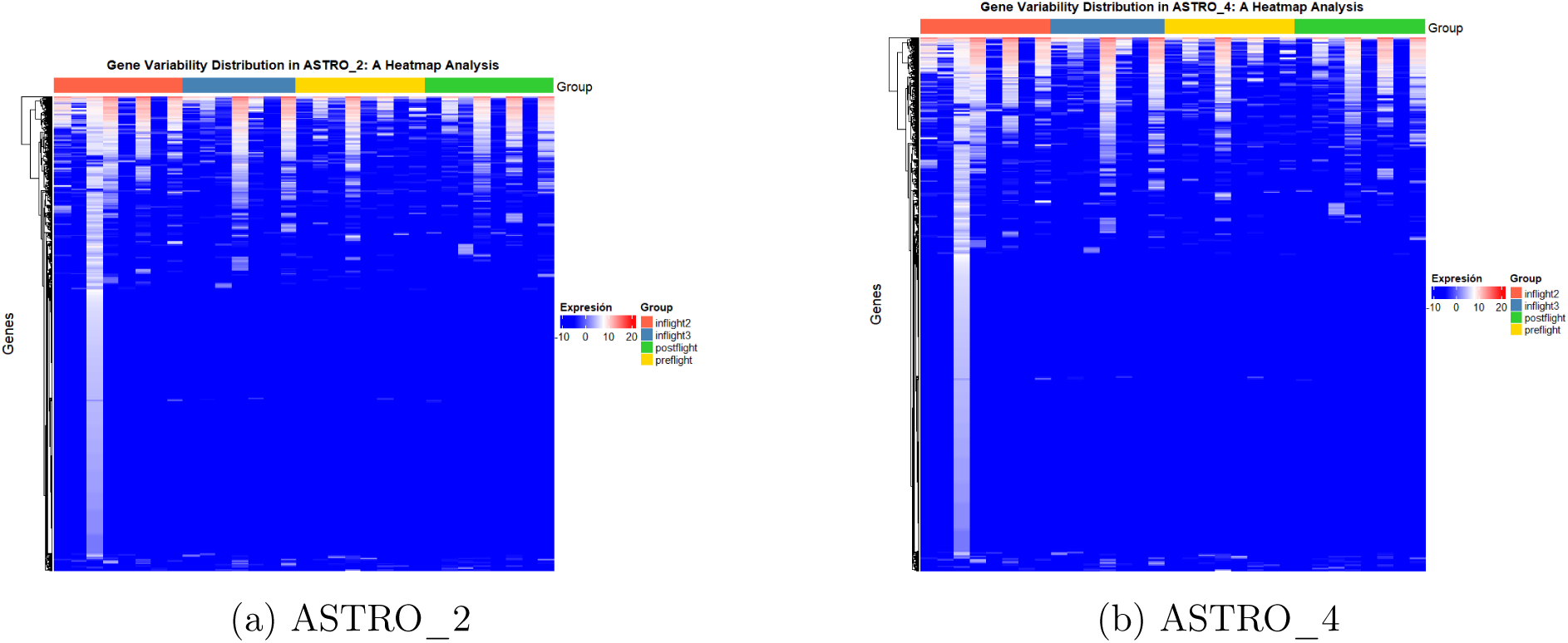
Heatmaps of most variable genes in scenarios ASTRO_2 and ASTRO_4.

**Figure 21:**
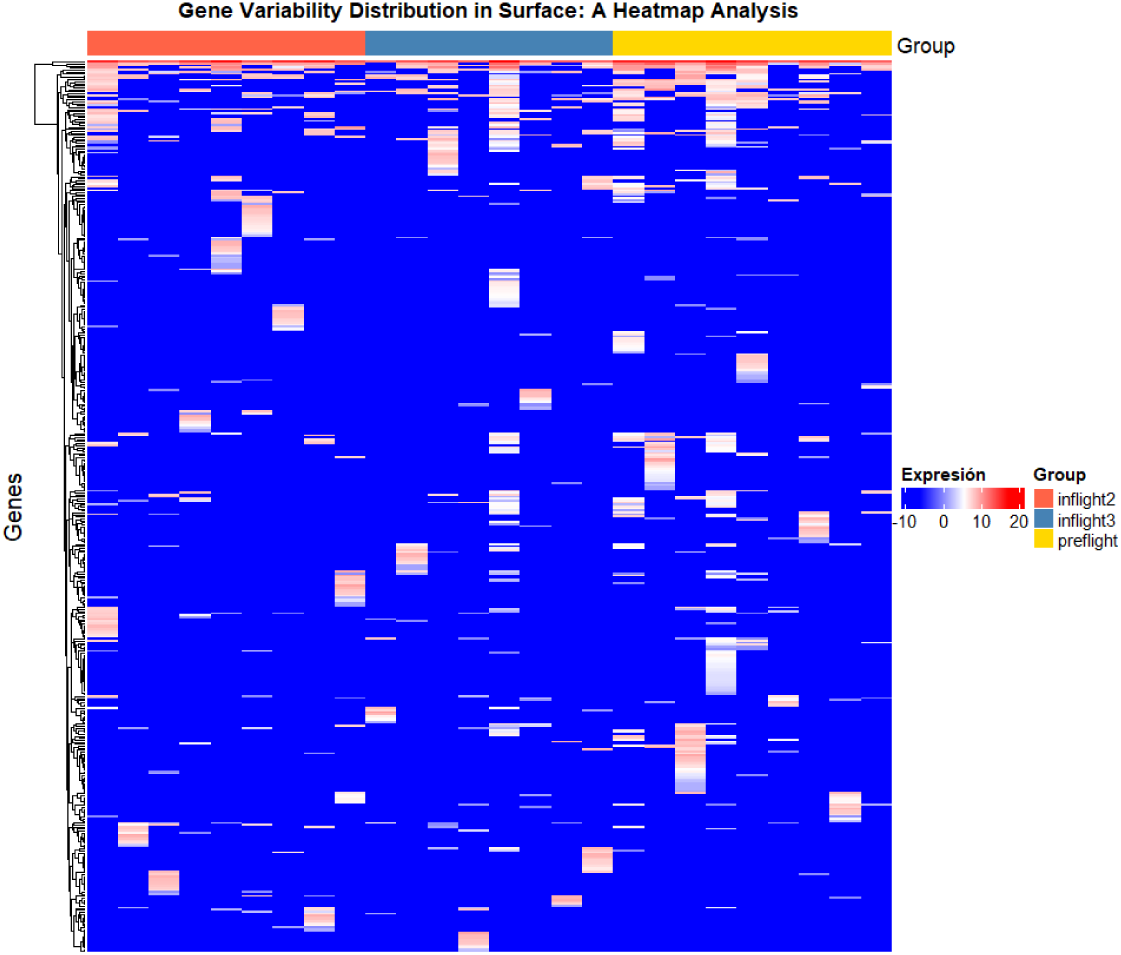
Heatmap of most variable genes in the Surface scenario.

Astro1 and Astro3 are the scenarios that show the greatest variability among the analyzed genes, evidenced by a greater presence of blank spaces and yellow tones in the heatmaps (see Figures 6b and 7b). In contrast, Astro2 and Astro4 present much less variable patterns, where blue ranges predominate indicating low expression, with only some discrete lines or gradients suggesting slight changes (see Figures 6c and 6d). The Surface scenario reflects the lowest global variability, characterized by an almost entirely blue heatmap (see Figure 7a). This behavior highlights the heterogeneity existing between the different scenarios. Additionally, it suggests that, in the case of the analyzed genes, these maintain considerably low expression values in the datasets associated with microbiota or surface. This could be due to competition with other microorganisms present in these samples, added to the fact that the performed analysis considers a general metatranscriptome, without focusing exclusively on Staphylococcus aureus expression.

## Appendix B Functional Enrichment

### Functional enrichment of modules after bootstrapping

After completing the *bootstrapping* analysis, functional enrichment of the modules that demonstrated the greatest robustness was performed. In microbiota 1 and microbiota 3, one robust module was obtained in each scenario, while in microbiota 2, microbiota 4, and BRIC_23, four robust modules were identified in each case. For the Surface scenario, two modules were detected, although it was not possible to perform a detailed functional association between genes and predominant functions for both modules simultaneously due to limitations in data availability.

The figures presented below illustrate the genes associated with the identified functions, the most predominant subcellular localization, and the most frequently detected functions.

**Figure 22:**
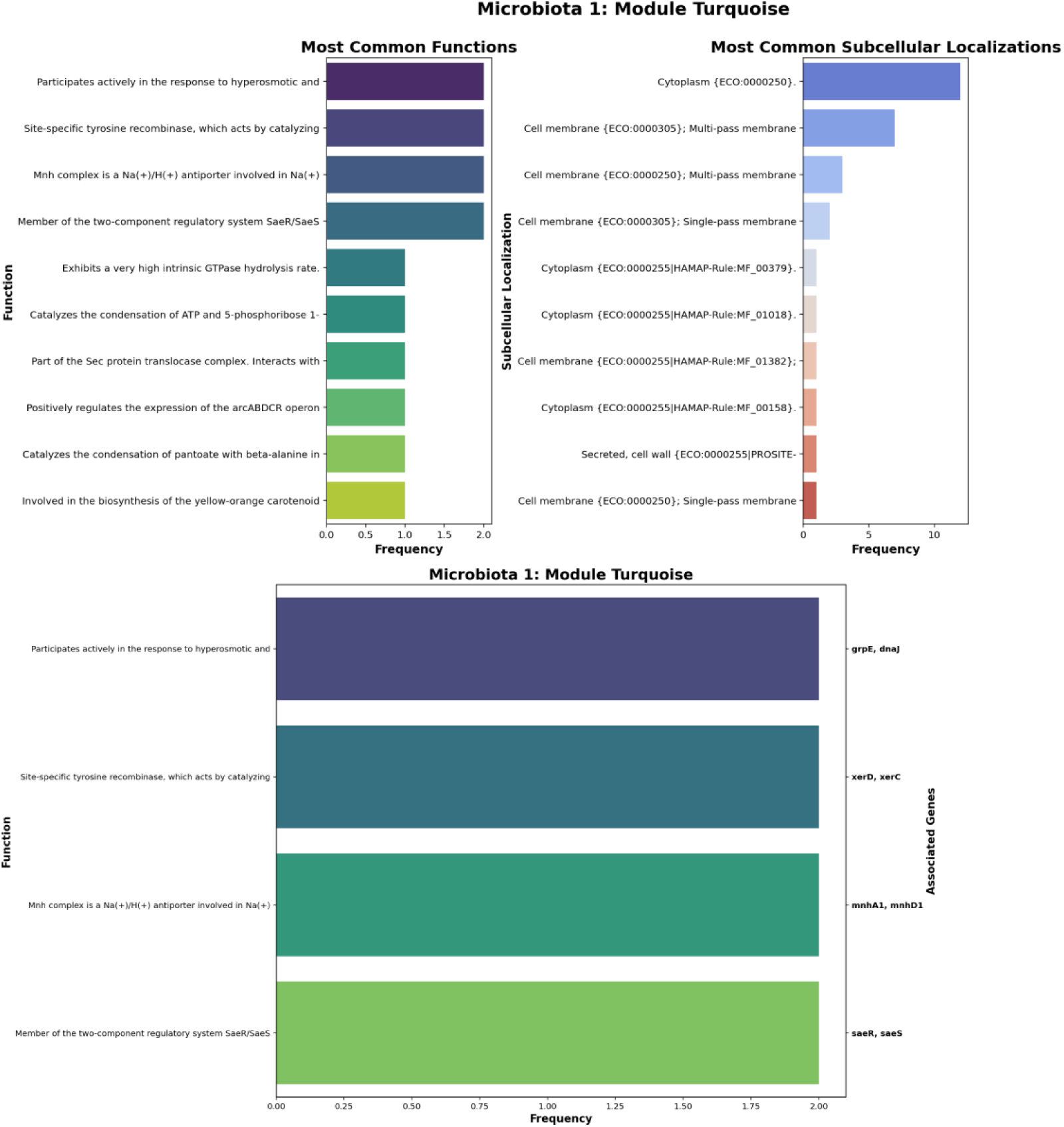
Representation of microbiota 1 scenario modules showing their genes along with their functions.

In the blue module of microbiota 1 (Figure 22), the *GrpE* and *DnaJ* genes play a crucial role in cellular stress response induced by thermal shock and hyperosmotic conditions. These proteins function coordinately with *DnaK* to prevent aggregation of denatured proteins, facilitating their proper folding and maintaining protein homeostasis. Additionally, *GrpE* acts as a nucleotide exchange factor, functioning as a thermosensor and modulating cellular response based on environmental temperature.

The *XerD* and *XerC* genes are responsible for chromosomal segregation during cell division through tyrosine-specific recombinases, in addition to maintaining segregational stability of plasmids. The complex encoded by *MnhA1/D1* regulates intracellular pH through a Na^+^/H^+^ antiporter, essential for bacterial survival in saline or alkaline environments. Finally, the SaeR/SaeS regulatory system controls the expression of virulence factors.

**Figure 23:**
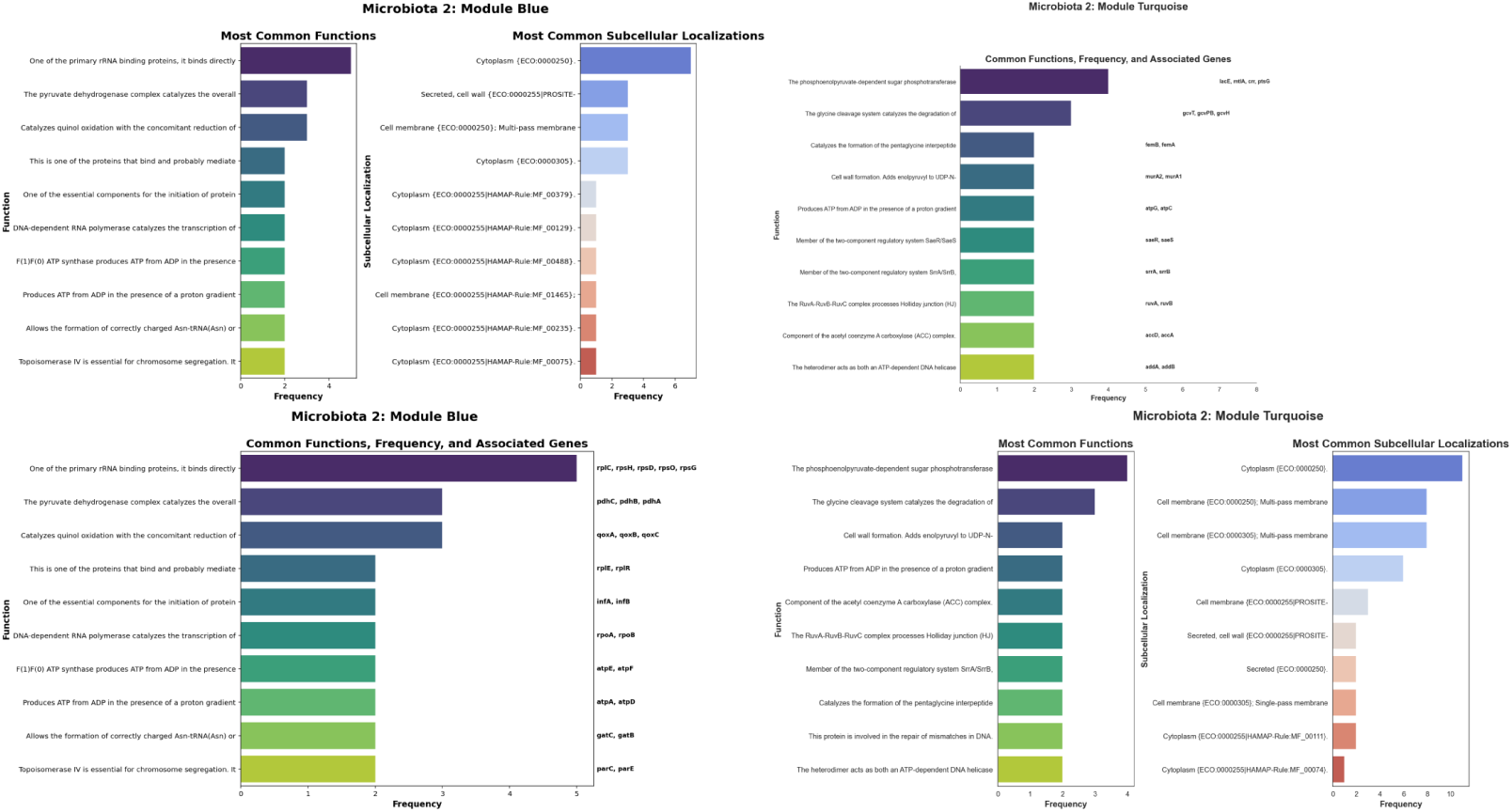
Representation of microbiota 2 scenario modules showing their genes along with their functions.

In microbiota 2 (Figure 23), the blue module includes genes such as *RplC* and *Rps*, essential for proper rRNA arrangement, as well as *Pdh*, which catalyzes the conversion of pyruvate to acetyl-CoA. In the turquoise module, the *crr/mtlA* (LACe) gene participates in the sugar phosphotransferase system (PTS), using PEP to transport and phosphorylate sugars. The predominant subcellular localizations are the cytoplasm and cell membrane.

**Figure 24:**
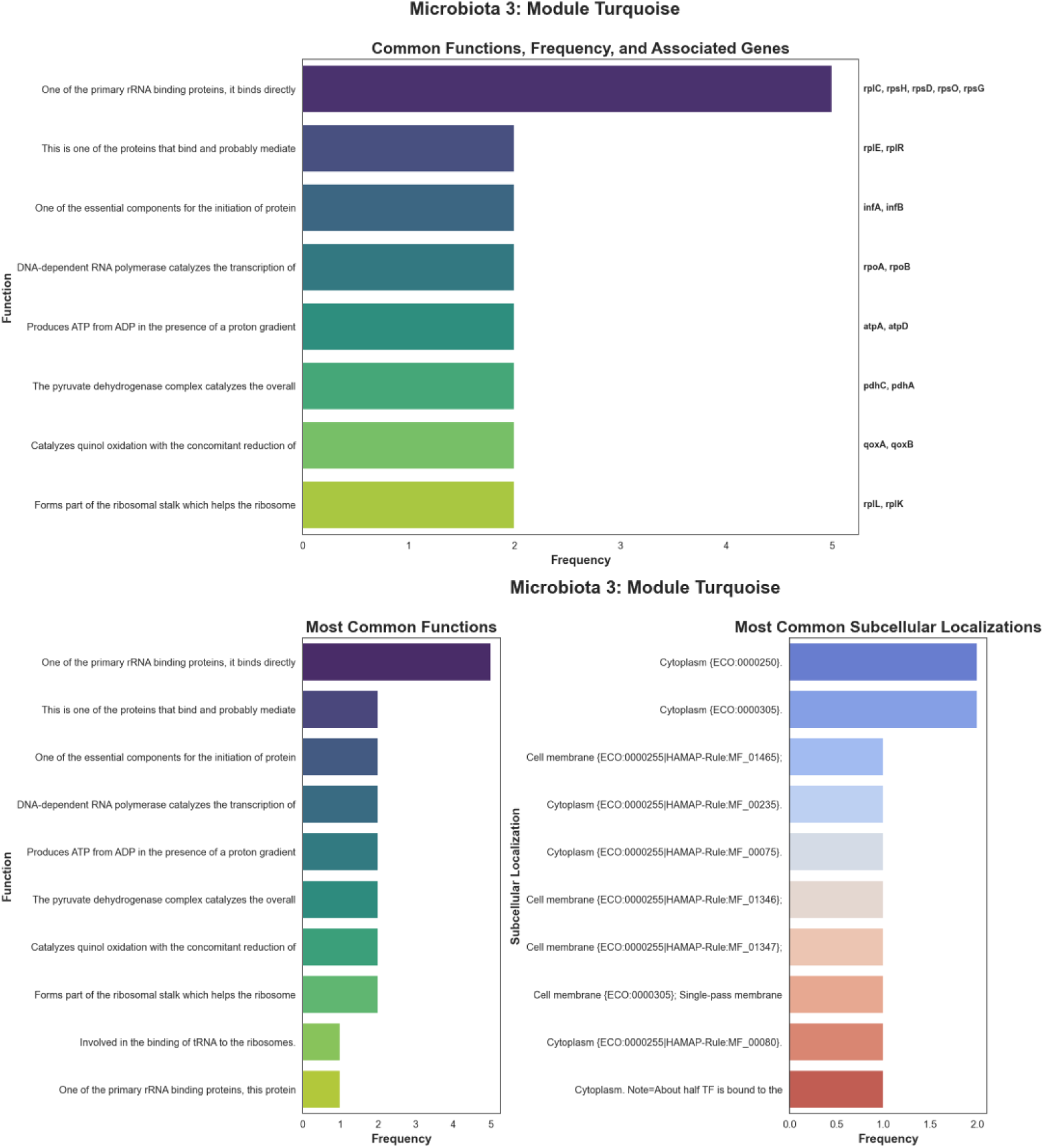
Representation of microbiota 3 scenario modules showing their genes along with their functions.

In microbiota 3 (Figure 24), the turquoise module also highlights genes such as *RplC*, *Rps*, and *Pdh*, reaffirming the relevance of ribosomal synthesis and energy metabolism. The majority subcellular localizations are the cytoplasm and membrane.

**Figure 25:**
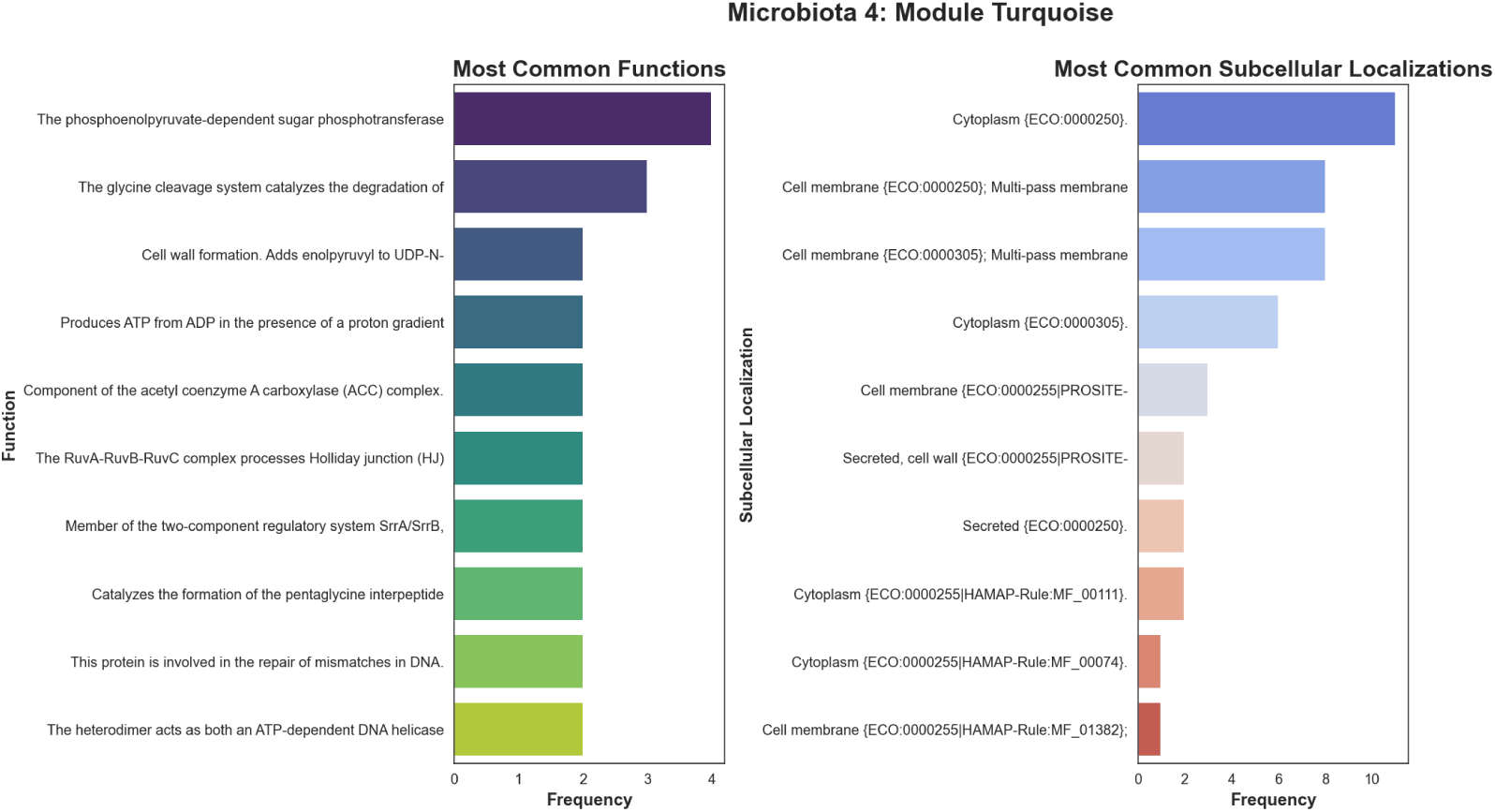
Representation of microbiota 4 scenario modules showing their genes along with their functions.

In microbiota 4 (Figure 25), the blue module concentrates genes involved in ribosomal synthesis and in the quinol oxidase complex (*qoxA*), essential for electron transfer. The turquoise module involves genes from the PTS system (*crr/mtlA*), the glycine cleavage system (*gcvT*), and peptidoglycan synthesis (*murA2*), key for maintaining cellular structural integrity.

**Figure 26:**
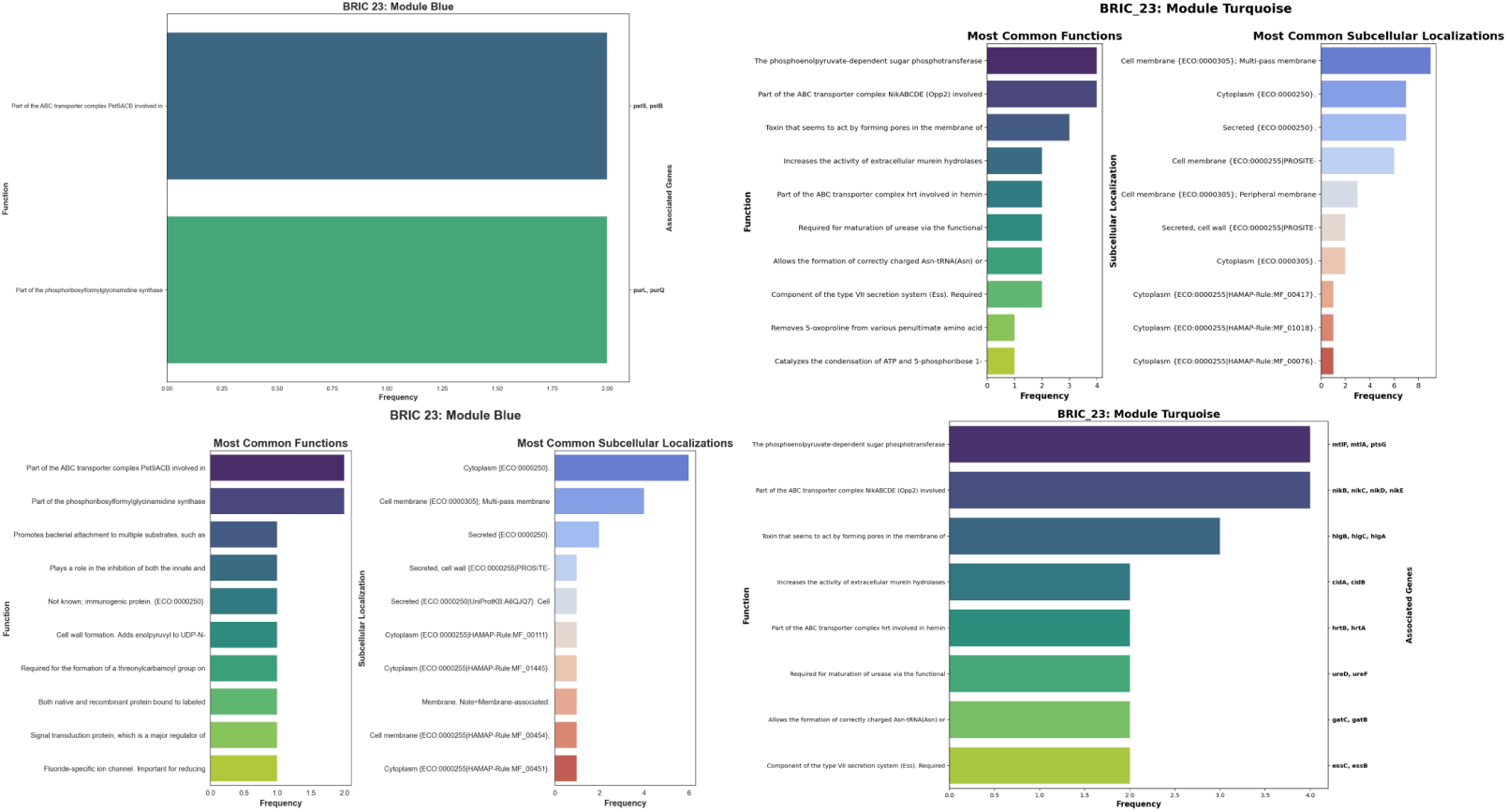
Representation of BRIC_23 scenario modules showing their genes along with their functions.

In BRIC_23 (Figure 26), the blue module stands out for genes related to ABC transport and ribosomal genes (*RplC*, *Rps*, *Pdh*), while the turquoise module involves the PTS system (*mtlF*), nickel transport (*nik*), and hemolytic toxins (*hlgA*, *hlgB*, *hlgC*), relevant for virulence and immune evasion.

**Figure 27:**
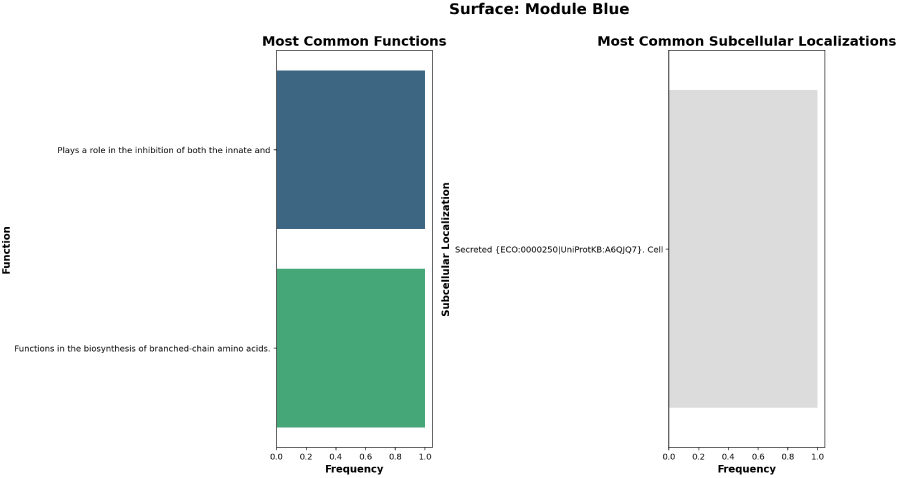
Representation of Surface scenario modules showing their genes along with their functions.

**Figure 28:**
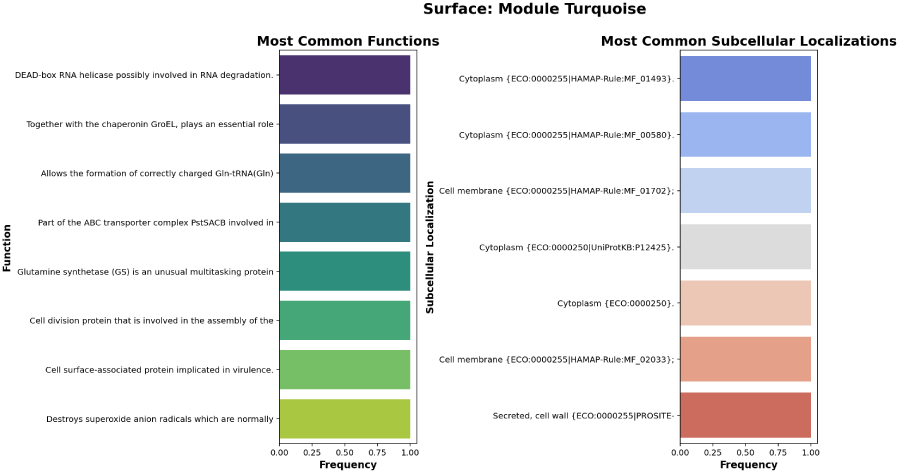
Representation of the turquoise module with surface genes along with their main functions

In the Surface scenario (Figure 27), the blue module includes the *Sbi* gene, which acts as a potent complement inhibitor, and *ilvD*, involved in branched-chain amino acid biosyn-thesis. In the turquoise module, *clfA* facilitates bacterial adhesion to human fibrinogen, while *sodM* eliminates reactive oxygen species, protecting the bacterial cell.

## Appendix C Genes of interest

The selected genes presented in Table 6 include a comprehensive list of *Staphylococcus aureus* loci analyzed in this study, along with their main functions, functional categories, and detailed descriptions. This table was generated using a longtable environment to accommodate a large number of entries spanning multiple pages, with increased row height (\renewcommand{\arraystretch}{1.2}) for improved readability. The genes are categorized based on key biological roles, including metabolism, gene regulation, stress response, protein synthesis, transport/membrane functions, secretion systems, virulence factors, adhesion, DNA replication, and essential cellular functions. Hypothetical proteins and uncharacterized loci are also listed to highlight areas of potential future research. This structured presentation allows readers to quickly identify genes of interest, understand their putative functions, and contextualize them within the broader transcriptomic and functional analyses discussed in this work.

**Table 6:** Selected genes with main functions and descriptions (Appendix C)

| **Gene** | **Main Function** | **Functional Category** | **Description** |
| --- | --- | --- | --- |
| SAR2748 | Hypothetical protein | Unknown function | Uncharacterized protein |
| SAR2749 | Hypothetical protein | Unknown function | Uncharacterized protein |
| SAR2750 | Hypothetical protein | Unknown function | Uncharacterized protein |
| SAR2009 | Putative transcription factor | Gene regulation | Potential transcriptional regulator |
| SAR2746 | Hypothetical protein | Unknown function | Uncharacterized protein |
| SAR2152 | Membrane protein | Transport/Membrane | Putative transmembrane transport protein |
| SAR2155 | Secretion system | Protein secretion | Secretion system component |
| SAR0625 | Ribosomal protein | Protein synthesis | Ribosomal machinery component |
| SAR2580 | Metabolic enzyme | Metabolism | Enzyme involved in metabolic pathways |
| SAR0832 | Virulence factor | Pathogenesis | Virulence-associated protein |
| SAR0842 | Cell wall protein | Cell structure | Cell wall component |
| SAR2709 | Transcriptional regulator | Gene regulation | Gene expression control |
| SAR2273 | Stress protein | Stress response | Response to adverse conditions |

| Gene | Main Function | Functional Category | Description |
| --- | --- | --- | --- |
| SAR1425 | ABC system | Transport | ABC transporter |
| SAR1424 | ABC system | Transport | ABC transporter (associated component) |
| SAR1026 | Metabolic enzyme | Metabolism | Central metabolism enzyme |
| SAR1284 | Adhesion protein | Adhesion/Virulence | Adhesin for host cell binding |
| SAR0965 | Sigma factor | Gene regulation | RNA polymerase sigma factor |
| SAR0398 | Virulence protein | Pathogenesis | Virulence factor |
| SAR0409 | Enzyme | Metabolism | Metabolic enzyme |
| SAR0009 | DnaA | DNA replication | Chromosomal replication initiator |
| SAR2126 | Hypothetical protein | Unknown function | Uncharacterized protein |
| SAR0490 | Membrane protein | Transport/Membrane | Transmembrane protein |
| SAR0506 | Enzyme | Metabolism | Metabolic enzyme |
| SAR1926 | Regulator | Gene regulation | Gene expression regulator |
| SAR2487 | Hypothetical protein | Unknown function | Uncharacterized protein |
| SAR0068 | Ribosomal protein | Protein synthesis | Ribosomal subunit |
| SAR2167 | Stress protein | Stress response | Heat shock protein |

| Gene | Main Function | Functional Category | Description |
| --- | --- | --- | --- |
| SAR2123 | Secretion system | Protein secretion | Type VII secretion system component |
| SAR2124 | Secretion system | Protein secretion | Type VII secretion system component |
| SAR2125 | Secretion system | Protein secretion | Type VII secretion system component |
| SAR0039 | Virulence protein | Pathogenesis | Virulence factor (duplicated entry) |
| SAR0966 | Sigma factor | Gene regulation | Alternative sigma factor |
| SAR0894 | Biosynthesis | Metabolism | Biosynthetic enzyme |
| SAR0895 | Biosynthesis | Metabolism | Biosynthetic enzyme (operon) |
| SAR0896 | Biosynthesis | Metabolism | Biosynthetic enzyme (operon) |
| SAR0897 | Biosynthesis | Metabolism | Biosynthetic enzyme (operon) |
| SAR0217 | Membrane protein | Transport/Membrane | Transmembrane protein |
| SAR0141 | Enzyme | Metabolism | Metabolic enzyme |
| SAR1187 | Regulator | Gene regulation | Transcriptional regulator |
| SAR1098 | Stress protein | Stress response | Stress response protein |
| SAR1772 | Enzyme | Metabolism | Amino acid metabolism enzyme |
| SAR2001 | Hypothetical protein | Unknown function | Uncharacterized protein |
| SAR1993 | Membrane protein | Transport/Membrane | Membrane protein |
| SAR0806 | Enzyme | Metabolism | Metabolic enzyme |

| Gene | Main Function | Functional Category | Description |
| --- | --- | --- | --- |
| SAR2713 | Regulator | Gene regulation | Expression regulator |
| SAR1974 | Stress protein | Stress response | Stress response protein |
| SAR0920 | Virulence factor | Pathogenesis | Virulence protein |
| SAR0721 | Enzyme | Metabolism | Nucleotide metabolism enzyme |
| SAR0060 | Ribosomal protein | Protein synthesis | Ribosomal protein |
| SAR0059 | Ribosomal protein | Protein synthesis | Ribosomal protein (adjacent to SAR0060) |
| SAR1831 | Membrane protein | Transport/Membrane | Transport protein |
| SAR0050 | Essential protein | Essential function | Essential protein for growth |
| SAR1211 | Enzyme | Metabolism | Metabolic enzyme |
| SAR2790 | Hypothetical protein | Unknown function | Uncharacterized protein |
| SAR1021 | Regulator | Gene regulation | Transcriptional regulator |
| SAR1142 | Stress protein | Stress response | Stress response protein |

## References

[1] Aurelie Crabbe, et al. “Response of Pseudomonas aeruginosa PAO1 to low shear modelled microgravity involves AlgU regulation”. In: Environmental Microbiology 12.4 (2010), pp. 1019–1028. doi: 10.1111/j.1462-2920.2010.02184.x. url: https://enviromicro-journals.onlinelibrary.wiley.com/doi/10.1111/j.1462-2920.2010.02184.x.

[2] Carina Audretsch, et al. “Modeling of stringent-response reflects nutrient stress induced growth impairment and essential amino acids in different Staphylococcus aureus mutants”. In: Scientific Reports 11.1 (2021), p. 9651. url: https://pubmed.ncbi.nlm.nih.gov/33958641/.

[3] Yoann Augagneur, et al. “Analysis of the CodY RNome reveals RsaD as a stressresponsive riboregulator of overflow metabolism in Staphylococcus aureus”. In: Molecular microbiology 113.2 (2020), pp. 309–325. url: https://pubmed.ncbi.nlm.nih.gov/31696578/.

[4] Bowtie: Manual. Accessed: 2025-06-10. 2025. url: https://bowtie-bio.sourceforge.net/manual.shtml.

[5] Kenny Zhi Ming Chen, Linda My Vu, and Amy Cheng Vollmer. “Cultivation in longterm simulated microgravity is detrimental to pyocyanin production and subsequent biofilm formation ability of *Pseudomonas aeruginosa*”. In: Microbiology Spectrum 12.10 (2024), e00211–24. doi: 10.1128/spectrum.00211-24. url: https://pmc.ncbi.nlm.nih.gov/articles/PMC11448113/.

[6] Thomas J Corydon, et al. “Current knowledge about the impact of microgravity on gene regulation”. In: Cells 12.7 (2023), p. 1043. url: https://pubmed.ncbi.nlm.nih.gov/37048115/.

[7] John G Decelle and Gerald R Taylor. “Autoflora in the upper respiratory tract of Apollo astronauts”. In: Applied and Environmental Microbiology 32.5 (1976), pp. 659–665. url: https://pubmed.ncbi.nlm.nih.gov/984836/.

[8] Centers for Disease Control and Prevention (CDC). Antimicrobial Resistance Threats in the United States, 2021–2022. url: https://www.cdc.gov/antimicrobial-resistance/data-research/threats/update-2022.html.

[9] Centers for Disease Control and Prevention (CDC). Methicillin-resistant Staphylococcus aureus (MRSA) is responsible for more than 70 000 severe infections and 9 000 deaths per year. url: https://www.dc.gov/mrsa/hcp/infection-control/index.html.

[10] Patricia Fajardo-Cavazos and Wayne L Nicholson. “Establishing standard protocols for bacterial culture in biological research in canisters (BRIC) hardware”. In: Gravitational and Space Research 4.2 (2016), pp. 58–69. url: https://www.researchgate.net/publication/343142041_Establishing_Standard_Protocols_for_Bacterial_Culture_in_Biological_Research_in_Canisters_BRIC_Hardware.

[11] Ákos Farkas and Gyula Farkas. “Effects of spaceflight on human skin”. In: Skin Pharmacology and Physiology 34.5 (2021), pp. 239–245. url: https://pubmed.ncbi.nlm.nih.gov/34058745/.

[12] Francine E Garrett-Bakelman, et al. “The NASA twins study: a multidimensional analysis of a year-long human spaceflight”. In: Science 364.6436 (2019), eaau8650. url: https://www.science.org/doi/10.1126/science.aau8650.

[13] Getting Started · kraken-ng/Kraken Wiki. Accessed: 2025-06-10. 2025. url: https://github.com/kraken-ng/Kraken/wiki/Getting-Started.

[14] Dimitri Golaz, et al. “RNA-seq analysis in simulated microgravity unveils down-regulation of the beta-rhizobial siderophore phymabactin”. In: npj Microgravity 10.1 (2024), pp. 1–8. url: https://www.nature.com/articles/s41526-024-00391-7.

[15] Harvard Medical School. Bootstrap Methods. https://www.hms.harvard.edu/bss/neuro/bornlab/nb204/statistics/bootstrap.pdf. Accessed: 2025-08-20. 2024. url: https://richardson.byu.edu/624/myl10/bootstrapping.pdf.

[16] Matthew R Hauserman, et al. “Altered quorum sensing and physiology of Staphylococcus aureus during spaceflight detected by multi-omics data analysis”. In: npj Microgravity 10.1 (2024). url: https://www.nature.com/articles/s41526-023-00343-7.

[17] How to Run FastQC from the Command Line. Accessed: 2025-06-10. 2025. url: https://olvtools.com/en/documents/fastqc.

[18] Hyang Jang, So Young Choi, and Robert J Mitchell. “Staphylococcus aureus sensitivity to membrane disrupting antibacterials is increased under microgravity”. In: Cells 12.14 (2023), p. 1907. url: https://www.mdpi.com/2073-4409/12/14/1907.

[19] KEGG. KEGG GENOME: Staphylococcus aureus subsp. aureus MRSA252 (MRSA). Accessed: 2025-06-10. 2025. url: https://www.kegg.jp/kegg-bin/show_organism?org=sar.

[20] W., et al. Kim. “Effects of spaceflight on Pseudomonas aeruginosa final cell density is modulated by nutrient and oxygen availability”. In: mBio 5.2 (2014), e01021–14. doi: 10.1128/mBio.01021-14. url: https://www.ncbi.nlm.nih.gov/pmc/articles/PMC4228280/.

[21] W., et al. Kim. “Spaceflight Promotes Biofilm Formation by Pseudomonas aeruginosa”. In: PLoS ONE 8.5 (2013), e62437. doi: 10.1371/journal.pone.0062437. url: https://journals.plos.org/plosone/article?id=10.1371/journal.pone.0062437.

[22] Bernhard Krismer, et al. “The commensal lifestyle of Staphylococcus aureus and its interactions with the nasal microbiota”. In: Nature Reviews Microbiology 15.11 (2017), pp. 675–687. url: https://pubmed.ncbi.nlm.nih.gov/29021598/.

[23] Peter Langfelder and Steve Horvath. “WGCNA: an R package for weighted correlation network analysis”. In: BMC bioinformatics 9.1 (2008), pp. 1–13. url: https://bmcbioinformatics.biomedcentral.com/articles/10.1186/1471-2105-9559.

[24] Shengkun Liu, et al. “The role of FTO in m6A RNA methylation and immune regulation in Staphylococcus aureus infection-related osteomyelitis”. In: Frontiers in Microbiology 16 (2025), p. 1526475. url: https://pubmed.ncbi.nlm.nih.gov/39980685/.

[25] Pedro Madrigal, et al. “Machine learning algorithm to characterize antimicrobial resistance associated with the International Space Station surface microbiome”. In: Microbiome 10.1 (2022), pp. 1–12. url: https://microbiomejournal.biomedcentral.com/articles/10.1186/s40168-022-01332-w.

[26] Paul R McAdam, et al. “Molecular tracing of the emergence, adaptation, and transmission of hospital-associated methicillin-resistant Staphylococcus aureus”. In: Proceedings of the National Academy of Sciences 109.23 (2012), pp. 9107–9112. url: https://www.pnas.org/doi/10.1073/pnas.1202869109.

[27] Arijit Mitra and Subhadeep Mukhopadhyay. “Regulation of biofilm formation by non-coding RNA in prokaryotes”. In: Current Research in Pharmacology and Drug Discovery 4 (2023). url: https://pubmed.ncbi.nlm.nih.gov/36636617/.

[28] Mark D Morrison and Wayne L Nicholson. “Meta-analysis of data from spaceflight transcriptome experiments does not support the idea of a common bacterial space-flight response”. In: Scientific reports 8.1 (2018), p. 13859. url: https://pubmed.ncbi.nlm.nih.gov/30258082/.

[29] NASA. Risk of Altered Immune System Responses. Accessed: 2025-04-23. 2025. url: https://www.nasa.gov/reference/risk-of-altered-immune-system-responses/.

[30] NASA Science. Polymicrobial Biofilm Growth and Control during Spaceflight (Bacterial Adhesion and Corrosion). Accessed: 2025-04-23. 2025. url: https://science.nasa.gov/biological-physical/investigations/bacterial-adhesion/.

[31] Eliah G Overbey, et al. “Collection of biospecimens from the inspiration4 mission establishes the standards for the space omics and medical atlas (SOMA)”. In: Nature Communications 15.1 (2024), pp. 1–14. url: https://www.nature.com/articles/s41467-024-48806-z.

[32] Eliah G Overbey, et al. “The space omics and medical atlas (SOMA) and international astronaut biobank”. In: Nature 632.8027 (2024), pp. 1145–1154. url: https://www.nature.com/articles/s41586-024-07639-y.

[33] Luciana Caenazzo Pamela Tozzo Arianna Delicati. “Skin microbial changes during space flights: a systematic review”. In: Life 12.10 (2022), p. 1498. url: https://www.mdpi.com/2075-1729/12/10/1498.

[34] Qi Peng, et al. “A review of biofilm formation of Staphylococcus aureus and its regulation mechanism”. In: Antibiotics 12.1 (2023), p. 12. url: https://pmc.ncbi.nlm.nih.gov/articles/PMC9854888/.

[35] Saugat Poudel, et al. “Coordination of CcpA and CodY regulators in Staphylococcus aureus USA300 strains”. In: mSystems 7.6 (2022), e00480–22. url: https://pubmed.ncbi.nlm.nih.gov/36321827/.

[36] Zahra Sedarat and Andrew W Taylor-Robinson. “Biofilm formation by pathogenic bacteria: applying a Staphylococcus aureus model to appraise potential targets for therapeutic intervention”. In: Pathogens 11.4 (2022), p. 388. url: https://pubmed.ncbi.nlm.nih.gov/35456063/.

[37] Jill Seladi-Schulman. MRSA survival rate by age, and factors that affect survival: 30-day. url: https://www.healthline.com/health/infection/mrsa-survival-rate-by-age.

[38] Wei Shi and Yang Liao. Rsubread/Subread Users Guide. 2025. url: https://bioconductor.org/packages/release/bioc/vignettes/Rsubread/inst/doc/SubreadUsersGuide.pdf.

[39] Xiaoyu Shi et al. “Identification of ferroptosis-related biomarkers for diagnosis and molecular classification of Staphylococcus aureus-induced osteomyelitis”. In: Journal of Inflammation Research 16 (2023), p. 1805. url: https://pubmed.ncbi.nlm.nih.gov/37131411/.

[40] Nitin Kumar Singh, et al. “Succession and persistence of microbial communities and antimicrobial resistance genes associated with International Space Station environmental surfaces”. In: Microbiome 6.1 (2018), pp. 1–23. url: https://microbiomejournal.biomedcentral.com/articles/10.1186/s40168-018-0585-2.

[41] Greg A Somerville and Richard A Proctor. “At the crossroads of bacterial metabolism and virulence factor synthesis in Staphylococci”. In: Microbiology and Molecular Biology Reviews 73.2 (2009), pp. 233–248. url: https://pubmed.ncbi.nlm.nih.gov/19487727/.

[42] Xin Su, et al. “Effects of simulated microgravity on the physiology of Stenotrophomonas maltophilia and multiomic analysis”. In: Frontiers in Microbiology 12 (2021), p. 701265. url: https://pubmed.ncbi.nlm.nih.gov/34512577/.

[43] L Tan, et al. “Therapeutic targeting of the Staphylococcus aureus accessory gene regulator (agr) system”. In: Frontiers in Microbiology 9 (2018), p. 322897. url: https://pubmed.ncbi.nlm.nih.gov/29422887/.

[44] Braden T Tierney, et al. “Longitudinal multi-omics analysis of host microbiome architecture and immune responses during short-term spaceflight”. In: Nature Microbiology 9.7 (2024), pp. 1661–1675. url: https://www.nature.com/articles/s41564-024-01635-8.

[45] UCLA Statistical Consulting Group. R Library: Introduction to Bootstrapping. https://stats.oarc.ucla.edu/r/library/r-library-introduction-to-bootstrapping/. Accessed: 2025-08-20. 2024. url: https://www.nature.com/articles/s41586-024-07639-y.

[46] USADELLAB. Trimmomatic: A flexible read trimming tool for Illumina NGS data. Accessed: 2025-06-10. 2025. url: https://pubmed.ncbi.nlm.nih.gov/24695404/.

[47] Bian Wang and Tom W Muir. “Regulation of virulence in Staphylococcus aureus: molecular mechanisms and remaining puzzles”. In: Cell chemical biology 23.2 (2016), pp. 214–224. url: https://pubmed.ncbi.nlm.nih.gov/26971873/.

[48] Virginia E Wotring. “Medication use by US crewmembers on the International Space Station”. In: The FASEB Journal 29.11 (2015), pp. 4417–4423. url: https://pubmed.ncbi.nlm.nih.gov/26187345/.

[49] Jie Zhou, et al. “Integrative omics analysis reveals insights into small colony variants of Staphylococcus aureus induced by sulfamethoxazole-trimethoprim”. In: BMC Microbiology 24.1 (2024), p. 212. url: https://pubmed.ncbi.nlm.nih.gov/38877418/.

